# Fc proteoforms of ACPA IgG discriminate autoimmune responses in plasma and synovial fluid of rheumatoid arthritis patients and associate with disease activity

**DOI:** 10.1101/2024.04.04.587930

**Authors:** Constantin Blöchl, Eva Maria Stork, Hans Ulrich Scherer, Rene E. M. Toes, Manfred Wuhrer, Elena Domínguez-Vega

## Abstract

Autoantibodies and their post-translational modifications (PTMs) are insightful markers of autoimmune diseases providing diagnostic and prognostic clues, thereby informing clinical decisions. However, current autoantibody analyses focus mostly on IgG1 glycosylation representing only a subpopulation of the actual IgG proteome. Here, by taking rheumatoid arthritis (RA) as prototypic autoimmune disease, we sought to circumvent these shortcomings and illuminate the importance of (auto)antibody proteoforms employing a novel comprehensive mass spectrometry (MS)-based analytical workflow. Profiling of anti-citrullinated protein antibodies (ACPA) IgG and total IgG in paired samples of plasma and synovial fluid revealed a clear distinction of autoantibodies from total IgG and between biofluids. This discrimination relied on comprehensive subclass-specific PTM profiles including previously neglected features such as IgG3 C_H_3 domain glycosylation, allotype ratios, and non-glycosylated IgG. Intriguingly, specific proteoforms were found to correlate with markers of inflammation and disease accentuating the need of such approaches in clinical investigations and calling for further mechanistic studies to comprehend the role of autoantibody proteoforms in defining autoimmune responses.

## Introduction

Autoantibodies directed against self-antigens are a hallmark of autoimmune diseases and important biomarkers for the clinics [1]. The role of autoantibodies in disease is manifold and ranges from direct pathogenic effects to being a consequence of prior inflammation and tissue damage [2, 3]. Rheumatoid arthritis (RA) is one of the most widespread autoimmune disease and serves as a prototypic autoimmune disease in many regards [4, 5]. RA is characterized by a systemic self-sustaining inflammation mainly of the synovial joints, which consequently results in a loss of cartilage and bone tissue [6, 7]. Pathogenesis of RA may span decades and is thought to be initiated by recognition of self-antigens associated with the production of autoantibodies [8]. Among those autoantibodies found in RA patients, anti-citrullinated protein antibodies (ACPA) are the most specific for the disease, present in about 70% of patients and generally identify a subset of RA patients with a worse prognosis [6, 9, 10]. While it is still under debate whether ACPA themselves directly contribute to the symptomatic hallmarks of RA [11, 12], ACPA are important for diagnosis, classification, prognosis, and potentially choice of treatment [13]. The ACPA response is evolving throughout the course of the disease and is associated with the onset of symptoms and the progression of RA [14]. The initial breach of tolerance and production of ACPA is followed by, e.g., epitope spreading, increasing isotype and subclass usage, and changes in glycosylation [15-20]. Regarding glycosylation, ACPA IgG adopt distinctive glycosylation hallmarks across Fab and Fc domains that have functional implications and inform about disease development [21-23]. Alterations in the Fc glycosylation of IgG in RA have been reported on different levels: (i.) total IgG glycans differ between RA patients and healthy individuals [24-28], (ii.) ACPA IgG glycans are different from total IgG [29-32], and (iii.) ACPA purified from synovial fluid (SF) differ in glycosylation from circulating ACPA [33]. Interestingly, ACPA Fc glycans change towards lower galactosylation before the onset of symptoms [19]. Low IgG galactosylation generally associates with inflammatory markers as well as disease activity in RA [26, 33, 34]. ACPA may include all four IgG subclasses with IgG1 and IgG4 being observed most frequently and at higher levels than the other subclasses [35, 36]. Together, previous studies thus emphasize the important role of ACPA for many aspects of RA. However, they are either focused on the determination of ACPA (subclass) titers based on enzyme-linked immunosorbent assays (ELISA) [35, 36] or IgG Fc glycosylation based on released glycan or glycopeptide mass spectrometric (MS) approaches [27, 33]. IgG subclasses and their sequence variants (allotypes), subclass-specific glycosylation, and other PTMs, cumulatively termed IgG proteoforms henceforth [37], are either not fully amenable to such approaches or neglected during analysis [19, 30]. Since these IgG features define interaction with downstream receptors and thus shape the immune response, comprehensive assessment of ACPA systemically and in affected tissues is critical to comprehend autoimmunity in RA and potentially predict disease activity or choice of treatment.

In this study, we leveraged the recent advancements in intact and subunit protein MS analysis to establish a comprehensive picture of the ACPA IgG Fc proteome in RA. Surpassing state-of-the-art glycopeptide analysis, we were able to assess subclass and allotype distributions, subclass-specific glycosylation, C_H_3 domain glycosylation, and PTM profiles for total and ACPA IgG from paired samples of plasma and SF of RA patients. Obtained IgG profiles portrayed ACPA responses in great detail visualizing the differences in ACPA characteristics compared to total and between ACPA IgG in plasma and SF. Several observed (auto)antibody proteoforms correlated with inflammatory markers and disease activity, emphasizing the need for comprehensive characterization of the ACPA response.

## Materials and methods

### Study design and patient details

Investigation of patient samples was approved by the medical ethical review board of the Leiden University Medical Center (code P17.151). All donors gave written informed consent for sample acquisition. Clinical characteristics of the investigated patients are listed in **Table S1**.

### Treatment of SF samples

To reduce viscosity, SF samples were treated by hyaluronidase (1 mg/ml; purified from bovine testicles, Sigma-Aldrich, St. Louis, MO, USA) at a ratio of 1:10 (v/v). Samples were vortexed rigorously for 3.0 min and incubated for 30.0 min at 37.0 °C while shaking. Samples were centrifuged for 10.0 min at 2000 xg to remove any potential precipitation.

### Capture of ACPA fractions

For ACPA capturing, neutravidin-coated beads (Pierce NeutrAvidin agarose; Thermo Fisher Scientific, Waltham, MA, USA) were freshly coupled with biotinylated CCP4 antigens (synthetized in-house [38]). Generally, 400-450 μl of beads were used per capturing experiment (depending on the numbers of sample) and initially washed four times with 1 ml phosphate-buffered saline (PBS; 0.035 mmol*/*L phosphate, 150 mmol*/*L NaCl, pH 7.6). Subsequently, the beads were incubated in PBS (equal volumes of beads and PBS) containing 2.0 μg of biotinylated CCP4 per 10 μl beads. This slurry was incubated for 1.0 h at 21 °C while shaking at 1,200 rpm on an Eppendorf shaker (Eppendorf, Hamburg, Germany). Excess of CCP4 peptide was removed by washing the beads four times with 1.0 ml PBS. The obtained CCP4-coupled beads were resuspended in a three-fold volume of PBS and loaded into 96-well filter plates (10 μm pore, Orochem Technologies, Naperville, IL, USA). 10 μl beads were added per well and washed thrice with 200 μl PBS followed each time by a centrifugation step (500 xg for 1.0 min). 25.0 μl of plasma and SF, respectively, were diluted to 200 μl with PBS and loaded to the washed beads. Of note, for patient 5 the amounts of plasma and SF were doubled due to the markedly lower ACPA IgG level (**Table S1**). Samples were incubated for 2.0 h at RT shaking at 900 rpm on an horizontal IKA shaker (IKA, Staufen, Germany). Centrifugation of the filter plates (500 xg, 2 min) yielded the non-ACPA containing flow-through that was stored at -20 °C until further processing. The retained beads were washed thrice with 200 μl PBS, followed by an incubation with 100 μl PBS at 37 °C on an Eppendorf shaker for 1.0 h to reduce non-specific binding of IgG. These beads were washed again thrice with 200 μl PBS and twice with 200 μl freshly prepared ammonium bicarbonate (50 mM; Sigma-Aldrich). After these washing steps, 2.5 U of IdeS (Genovis, Lund, Sweden) were added in 20 μl of 50 mM ammonium bicarbonate to the beads. The plate was incubated for 3.0 h in a moisture box at 37 °C without shaking. Fc/2 fractions were recovered by centrifugation at 500 xg for 2.0 min and directly measured by nanoscale HPLC-MS as specified below. Specificity of the capturing was assessed by conducting this workflow with three ACPA-negative plasma and three ACPA-negative SF samples.

### Determination of ACPA IgG levels

ACPA IgG levels of RA plasma and SF samples were determined using ELISA. Briefly, streptavidin-coated plates (standard capacity; Microcoat Biotechnologie GmbH, Germany) were incubated for one hour at room temperature with 1.0 μg/ml N-terminally biotinylated CCP2 (patent number: EP2071335) diluted in PBS supplemented with 0.1% bovine serum albumin (BSA). Samples were diluted in PBS with 1% BSA and 0.05% Tween (PBT) and incubated for 1 h at 37 °C. Bound ACPA IgG was detected by incubation with horseradish peroxidase-labelled rabbit anti-human IgG (P0214; Agilent Dako, CA, USA) diluted 1:8000 in PBT for 1 h at 37 °C before the ELISA was read out using ABTS and H_2_O_2_ at an iMark Microplate Absorbance Reader (Bio-Rad Laboratories, Inc., CA, USA). ACPA IgG levels were quantified based on an in-house standard of pooled RA patient plasma using the Microplate manager software MPM-6 (Bio-Rad Laboratories, Inc.). Unspecific reactivity to the peptide backbone was assessed by applying CArgP2, the non-modified version of CCP2, instead of CCP2.

### Capture of total IgG fractions

Total IgG fractions were purified from the flow-through obtained after ACPA purification of plasma and SF samples. The procedure to capture total IgG by affinity purification was described in detail in Blochl et al. [39].

Briefly, diluted plasma or SF samples corresponding to approx. 8 μg IgG (considering a concentration of 10 μg*/*μl IgG in both biological fluids [40]) were loaded onto Fc-specific beads (CaptureSelect (FcXL); Thermo Fisher Scientific) that were packed into 96-well filter plates. A 2.0 h incubation step was followed by washing multiple times with PBS and subsequent on-bead digestion with IdeS, which cleaves IgG below the hinge region. Released Fab_2_ portions were washed from the beads, followed by release of Fc subunits under acidic conditions (0.1 mol/L formic acid). Eluted Fc subunits were directly subjected to nanoscale reversed phase (RP) HPLC-MS analysis without any prior purification steps.

### Deglycosylation of IgG C_H_2 domain glycans

25 μl plasma were diluted to 200 μl with PBS and deglycosylated at the C_H_2 domain glycosylation site by EndoS2 (Glycinator; Genovis) at a ratio of 1 U/μg IgG for 2.5 h at 37 °C. Subsequently, ACPA and total IgG Fc/2 portions were purified from these C_H_2 domain-deglycosylated samples as described above.

Monoclonal IgG3 allotypes (IGHG3*01 and IGHG3*11) were deglycosylated with EndoS2 (1 U/μg IgG for 2.5 h at 37 °C) followed by IdeS cleavage (2 U/μg IgG in 50 mM ammonium bicarbonate) and subsequently characterized by nano RP HPLC-MS. 200 ng of monoclonal IgG3 digests were injected in triplicates. Production and purification of these monoclonals is described in de Taeye et al. [41].

### Assessment of Fc-based capturing of IgG3 allotypes

Monoclonal IgG3 allotypes (IGHG3*01 and IGHG3*11) were directly digested by IdeS (2 U/μg IgG in 50 mM ammonium bicarbonate) or during FcXL-based capturing as described above. Obtained Fc/2 portions were measured by the designated nano RP HPLC-MS approach. 200 ng of monoclonal IgG3 digests were injected in triplicates.

### Assessment of reduced disulfide bonds

Deamidation of a monoclonal IgG1 (Herceptin; Roche Diagnostics, Penzberg, Germany) was induced by incubating the antibody at a concentration of 1.0 mg/ml in 200 mM ammonium bicarbonate (pH 8.4) at 37 °C for 7 days. Deamidated and non-deamidated control samples were cleaved by IdeS at a concentration of 2 U/μg IgG during incubation for 1 h at 37 °C and subjected to analysis. For the generation of reduced species, Fc/2 portions obtained from native Herceptin were diluted to 0.1 mg/ml and a final concentration of 4 M guanidinium HCl (Thermo Fisher Scientific) supplemented with 25 mM TCEP and were incubated at 60 °C for 5 min. For alkylation, Fc/2 portions obtained from Herceptin (0.1 mg/ml) were denatured in 4 M guanidinium HCl at 60 °C for 5 min. The sample was cooled down to 21 °C and alkylated by the addition of iodoacetamide (Sigma-Aldrich) to 25 mM followed by incubation at 21 °C for 30 min in the dark. Samples were immediately subjected to nano RP HPLC-MS analysis.

### High-performance liquid chromatography coupled to mass spectrometry (HPLC-MS)

Nano RP HPLC-MS measurements of intact Fc/2 subunits were relying on a recently developed approach [39]: Measurements were conducted on an UltiMate 3000 nanoRSLC system (Thermo Fisher Scientific) equipped with a C4 trap column (5 × 0.3 mm i.d., Acclaim™ PepMap™, 300 Å pore size; Thermo Fisher Scientific) and a diphenyl reversed phase column (150 × 0.1 mm i.d, Halo Bioclass, 1000 Å pore size; Advanced Material Technology, Wilmington, DE, USA). The exact conditions of separation are listed in Blochl et al. [39]. For analysis of ACPA Fc/2 subunits purified from plasma and SF, 5.0 μl of sample were injected. For total IgG, 0.75 μl of sample were injected. The nano RP HPLC system was hyphenated to a qTOF mass spectrometer (maXis Impact HD, Bruker Daltonics, Bremen, Germany) employing a nano ESI source (CaptiveSpray). Acetonitrile was used as dopant to enrich the nitrogen gas aiding ionization. In general, all measurements were conducted as technical duplicates. System performance was assessed by an Fc/2 standard obtained from a commercial mAb preparation (Herceptin; Roche Diagnostics).

### Data evaluation

Allotype constitutions of investigated patients were determined by intact mass matching after deconvolution in the Compass DataAnalysis software (Maximum Entropy deconvolution, Data point spacing at 1.0, MS resolution of 5,000; Bruker). Masses were picked based on the Sum Peak algorithm embedded in Compass DataAnalysis. For quantification of allotype-related glyco- and proteoforms, acquired MS data was pre-processed: MS files were converted into the .mzxml format sub-setting *m/z* (*m/z* 1000-1600) and retention time range (600-1320 s) employing msConvert [42]. Processed files were subjected to Gaussian smoothing of individual mass spectra (default settings, peak width of *m/z* 1.0) and subsequent baseline subtraction within these individual spectra with the Tophat algorithm (default settings, width *m/z* 1.0) within the OpenMS environment [43]. Prior to retention time alignment, data files were grouped into several packages based on the highest overlap in allotype constitution. In essence, files belonging to the same patient and patients with overlapping allotype profiles were grouped to allow for accelerated quantification. Retention time alignment was accomplished with Lacy tools [44]. These processed files were quantified in Skyline (22.2.0.351) [45]. As reported previously in detail [39], three charge states were considered for generation of extracted ion chromatograms (EICs) that were integrated for determination of peak areas. EIC *m/z*-borders were automatically calculated by Skyline considering the base peak isotope of individual analytes and a TOF resolution of 15,000. Analytes were integrated automatically and curated manually considering retention time and mass errors. In many instances, the synchronized integration feature embedded in Skyline could be used as the data featured aligned retention times. Skyline reports were further processed within the R environment [46]. Glycoforms of IgG3 allotypes that show occupancy of both *N*-glycosylation sites were relatively quantified after deconvolution in the Compass DataAnalysis software (settings as stated above). Site occupancy of the C_H_3 domain glycosylation site in IgG3 was determined by quantification of the doubly glycosylated H5N2, H3N4F1 glycoform within Skyline. The obtained peak area of this glycoform was divided by its relative abundance among all doubly glycosylated glycoforms as obtained from deconvolution, resulting in the total peak area of all glycoforms of doubly glycosylated IgG3. Comparison of this peak area with the combined peak area of the singly glycosylated IgG3 resulted in the site occupancy of the C_H_3 domain.

Identified and monitored C_H_2 domain IgG glycans are listed in **Table S2** including the used nomenclature. Calculation of glycosylation traits was based on the summation of weighted relative abundances of respective glycoforms as described in detail previously [47]. Integration of C_H_2 and C_H_3 domain glycans to annotate doubly glycosylated IgG3 was conducted employing MoFi [48]. Data visualization and statistical assessment was conducted in GraphPad Prism 9.3.1 and in R [46]. Allotype nomenclature was based on IMGT accession numbers [49, 50] that is linked to the Gm nomenclature as detailed previously [39, 51].

### Data availability

The raw mass spectrometry data is available under: https://doi.org/10.5281/zenodo.10261169.

## Results

### Subclasses and their glycosylation discriminate total and ACPA IgG from plasma and SF

To comprehensively characterize IgG Fc characteristics of RA patients, we expanded on a recently developed workflow to purify and analyze intact Fc subunits of ACPA and total IgG fractions from paired plasma and SF samples [39]. In brief, ACPA and total IgG were purified by antigen-or Fc-specific affinity beads, respectively followed by the release of intact single chain Fc subunits (Fc/2) by proteolysis with the protease IdeS (**Figure 1a**). A detailed visualization of both capturings is provided in **Figure S1a**. Fc/2 portions obtained from IgG fractions, i.e., total IgG from plasma, total IgG from SF, ACPA IgG from plasma, and ACPA IgG from SF, were characterized by a recently developed nanoscale RP HPLC-MS approach [39]. Eleven RA patients were investigated, whose characteristics are summarized in **Table S1**. Specificity of the ACPA capturing approach was assessed by including three plasma and three SF samples obtained from ACPA-negative patients (**Figure S1b**). ACPA levels assessed by nano RP HPLC-MS correlated well with ACPA IgG levels obtained by ELISA (**Figure S1c**). Precision of these measurements was assessed for a subset of three RA patients regarding total ACPA IgG signals (**Figure S2a**) as well as total/ACPA IgG allotype distributions (**Figure S2b)** and glycosylation patterns of IgG1 and IgG4 of ACPA (**Figure S3**) and total IgG (**Figure S4**).

**Figure 1.**
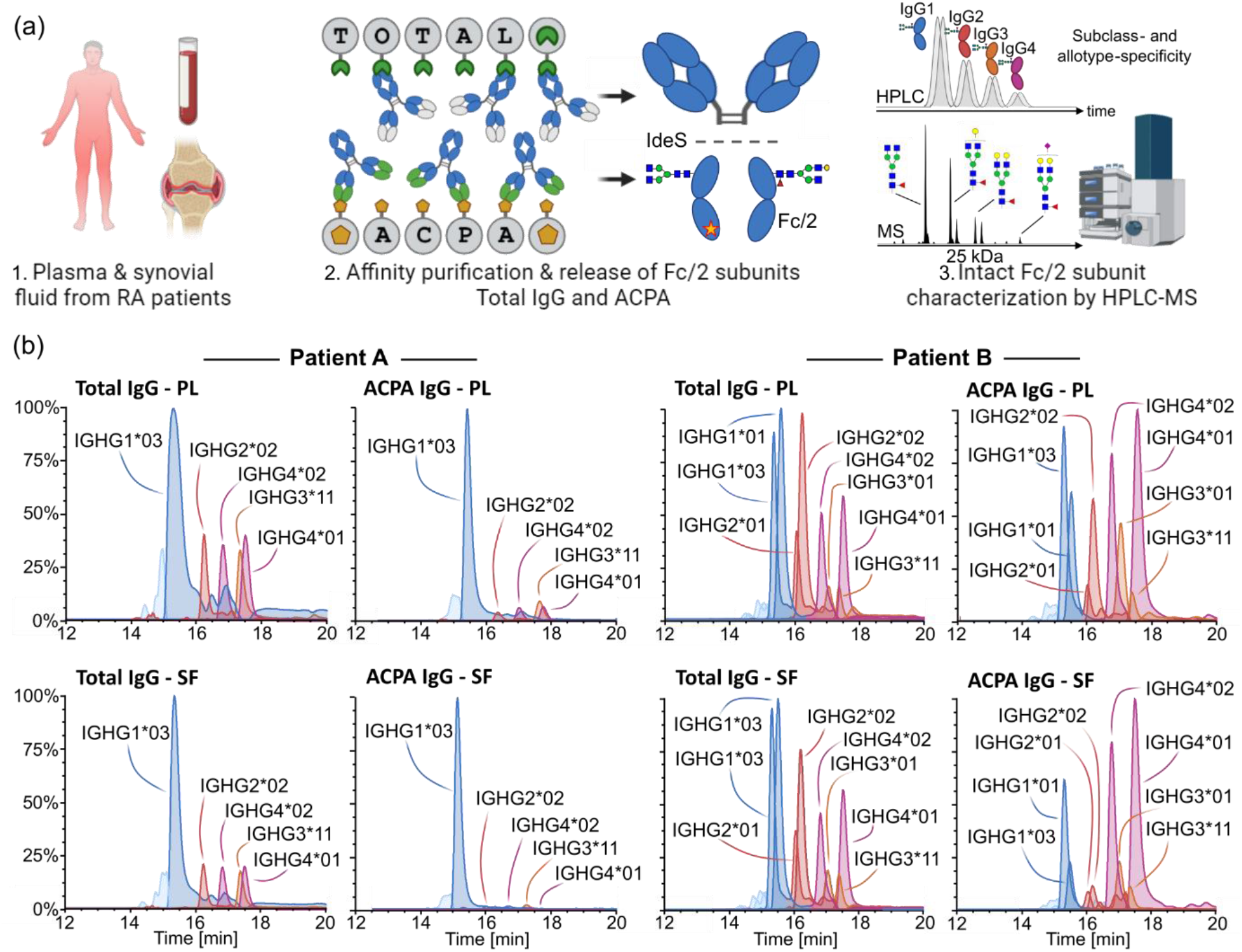
Schematic overview of the IgG (auto)antibody characterization workflow and Fc profiles of two exemplary patients. (a) Paired plasma and SF samples were analyzed from 11 RA patients. ACPA IgG and total IgG fractions were obtained by means of affinity purification employing cyclic citrullinated antigen-coated and Fc-specific beads, respectively. Fc/2 portions were released by specific proteolysis in the hinge region using IdeS and subjected to nanoscale HPLC-MS analysis. (b) Exemplary Fc profiles obtained from patient A (#3) and B (#2) for total and ACPA IgG purified from plasma and SF. Each allotype is represented by an EIC trace and color-coded by subclass (IgG1: blue, IgG2: red, IgG3: orange, and IgG4: violet). Relative abundances are shown in percent on the y-axis. Notably, as these EICs were generated based on G0F glycoforms (composition H3N4F1), differences in glycosylation may bias relative abundances of depicted allotypes.

The applied approach is sensitive for alterations in the Fc portion and thus resolves IgG sequence variations referred to as allotypes. Since each individual may express up to two allotypes per IgG subclass, a maximum of eight allotypes may be identified per patient. As exemplified in **Figure 1b**, ACPA and total IgG allotype abundances were highly variable between patients: Patient A comprised five allotypes and showed the common distribution of subclasses in total IgG in plasma and SF, i.e., high IgG1 abundances and lower amounts of IgG2, 3, and 4. In contrast, both ACPA profiles from this patient virtually lacked IgG2, 3, and 4 and were dominated by IgG1. Patient B comprised eight allotypes and showed next to IgG1 high abundance of IgG2 and IgG4 allotypes in total IgG fractions. Of note, ACPA profiles of this patient showed even higher levels of IgG4 allotypes, which was most pronounced in ACPA derived from SF compared to plasma.

Next to allotype and subclass abundance, the intact Fc subunit analysis strategy provided information on all coexisting Fc structural characteristics. Initially, subclass-specific glycosylation was assessed and summarized as specific glycosylation traits. An unsupervised principal component analysis was used to visualize differences in these glycosylation traits, e.g., galactosylation or sialylation, as well as relative subclass abundances (**Figure 2a**). In the score plot, a separation of total and ACPA IgG was evident. This was mainly driven by lower levels of afucosylation, bisection, and non-glycosylated IgG in ACPA purified from both plasma and SF as evident from the loading plot of the PCA. Remarkably, these differences seemed to be largely independent of subclass with the exception of IgG4 afucosylation and non-glycosylated IgG4 that were largely unaltered between ACPA and total IgG (**Figure S5** and **S6**). Regarding subclasses, ACPA show enhanced levels of IgG1 and IgG4 compared to total IgG, whereas IgG2 and IgG3 were generally low in ACPA (**Figure 2a, Table S3**). No obvious differences between ACPA IgG from plasma and SF were found regarding subclass usage. Moreover, a separation between plasma and SF IgG was observable. This was clearly more pronounced in ACPA and was mainly driven by differences in galactosylation and sialylation that were higher in ACPA from plasma. Significant differences between total and ACPA IgG were observed for IgG1 fucosylation (**Figure 2b**), bisection, and non-glycosylated IgG1 in plasma (**Figure 2c**) and SF (**Figure 2d**). Differences between total and ACPA IgG glycosylation traits in other subclasses were visualized in **Figure S5** and **S6** (data provided in **Table S3)**. The most pronounced differences in glycosylation between ACPA from plasma and SF, i.e., (a)galactosylation and sialylation, were visualized for IgG1 (**Figure 2e** and **f**) and IgG4 (**Figure 2g**). A complete comparison of glycosylation traits between ACPA from plasma and SF was compiled for **Figure S5** and **S6** (data in **Table S3**).

**Figure 2.**
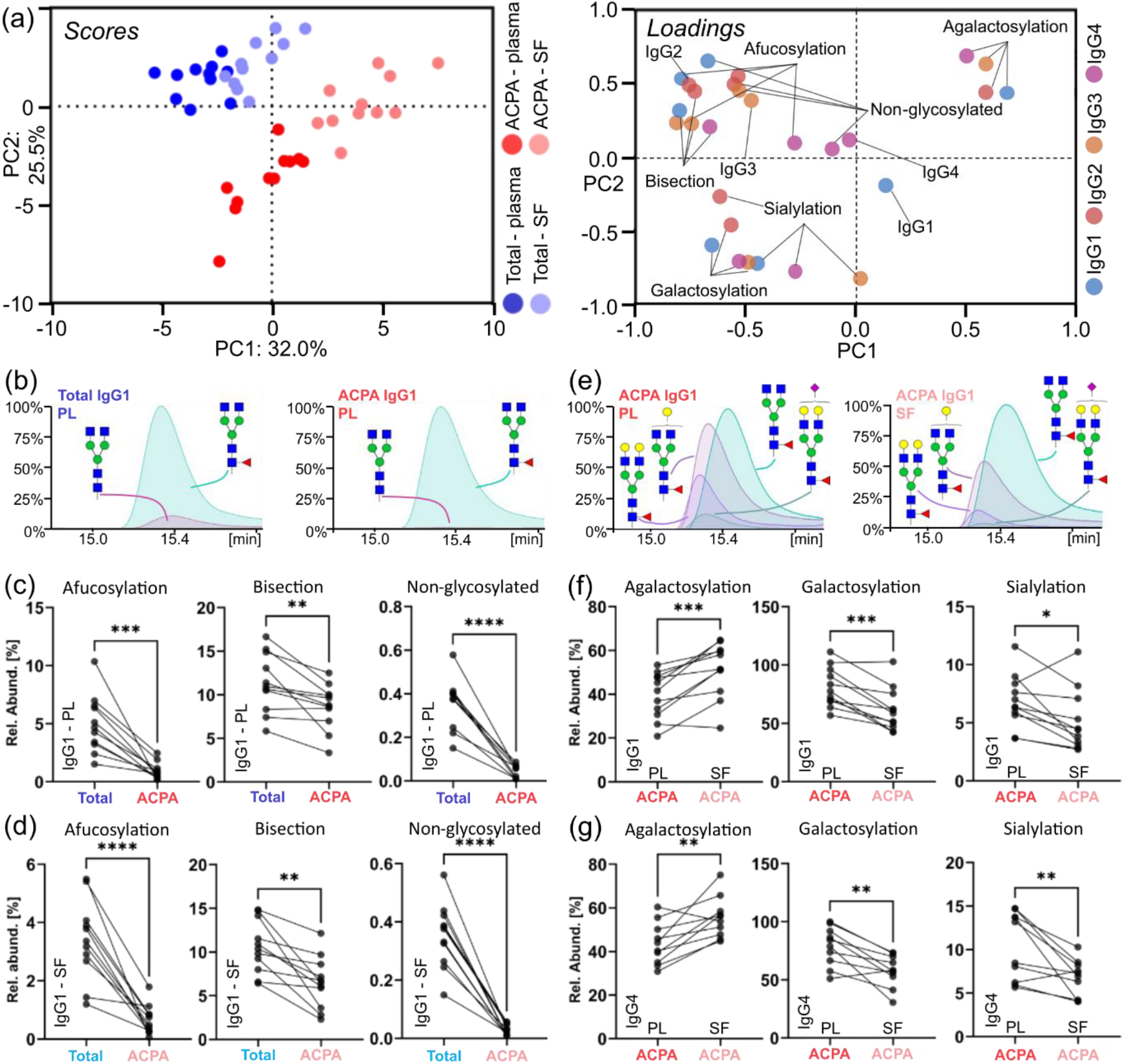
IgG subclasses and their glycosylation define ACPA and total IgG fractions in plasma (PL) and SF. (a) Principal component analysis integrating data on IgG subclass abundances, and subclass-specific glycosylation in the form of derived traits. Differences of IgG features in the four fractions within each patient were used as input. In the left panel, the score plot depicts data points representing individual patients. Each patient is represented by a set of four circles (blue: total IgG from plasma; light blue: total IgG from SF; red: ACPA IgG from plasma; light red: ACPA IgG from SF). In the right panel the corresponding loading plot is shown. Variables are colored by their association with one of the four subclasses. (b) Exemplary combined EIC of two glycoforms of the IGHG1*03 allotype visualizing the differences in afucosylation between total IgG (left) and ACPA IgG (right) in plasma. Differences in derived traits comparing total and ACPA IgG purified from (c) plasma or (d) SF. (e) Exemplary combined EIC of the major glycoforms of the IGHG1*03 allotype demonstrating the differences in galactosylation and sialylation levels comparing ACPA purified from plasma and SF, respectively. Differences in derived traits of ACPA (f) IgG1 and (g) IgG4 comparing plasma and SF. Significance levels were assessed by paired t-tests (* for p ≤ 0.05, ** for p ≤ 0.01, *** for p ≤ 0.001, and **** for p ≤ 0.0001).

### Allotype profiling reveals altered expression ratios between total and ACPA IgG

Analysis of intact Fc/2 subunits enabled monitoring of allotype constitution and expression. In **Figure 3a**, the allotype constitution of the 11 investigated RA patients is depicted showcasing the high variability of allotypes found even in this relatively small cohort. Intriguingly, the expression of allotypes markedly differed by the specific allotype sequence. For instance, the IGHG2*01/02 ratio was assessed in five applicable patients that showed this heterozygosity (**Figure 3b**). Total IGHG2*01 was lower expressed in all instances besides a high variability in this ratio between patients. Unequal expression was additionally found for patients bearing the common IGHG1*01/03 or IGHG4*01/02 allotype heterozygosity. Altered expression ratios of allotypes belonging to the same subclass were also assessed for ACPA. In **Figure 3c**, the IGHG1*01/03 ratio in total IgG (SF) and ACPA IgG (SF) is exemplarily depicted as EICs for one patient. Although for this patient these allotypes were virtually equally expressed in total IgG, ACPA showed a strong reduction in IGHG1*01 usage. These alterations in IgG1 allotype usage was found to be significant for both plasma and SF and were consistent for all applicable patients (**Figure 3d**). Likewise, IgG2 (**Figure 3e**) and IgG4 allotypes (**Figure 3f**) were investigated: Although variations in the IGHG2*01/02 and IGHG4*01/02 ratios were detected, no clear trend between total IgG and ACPA was observable. IgG3 allotype ratios could not be assessed due to the low amount of patients (≤ 2) that shared the same two IgG3 allotypes.

**Figure 3.**
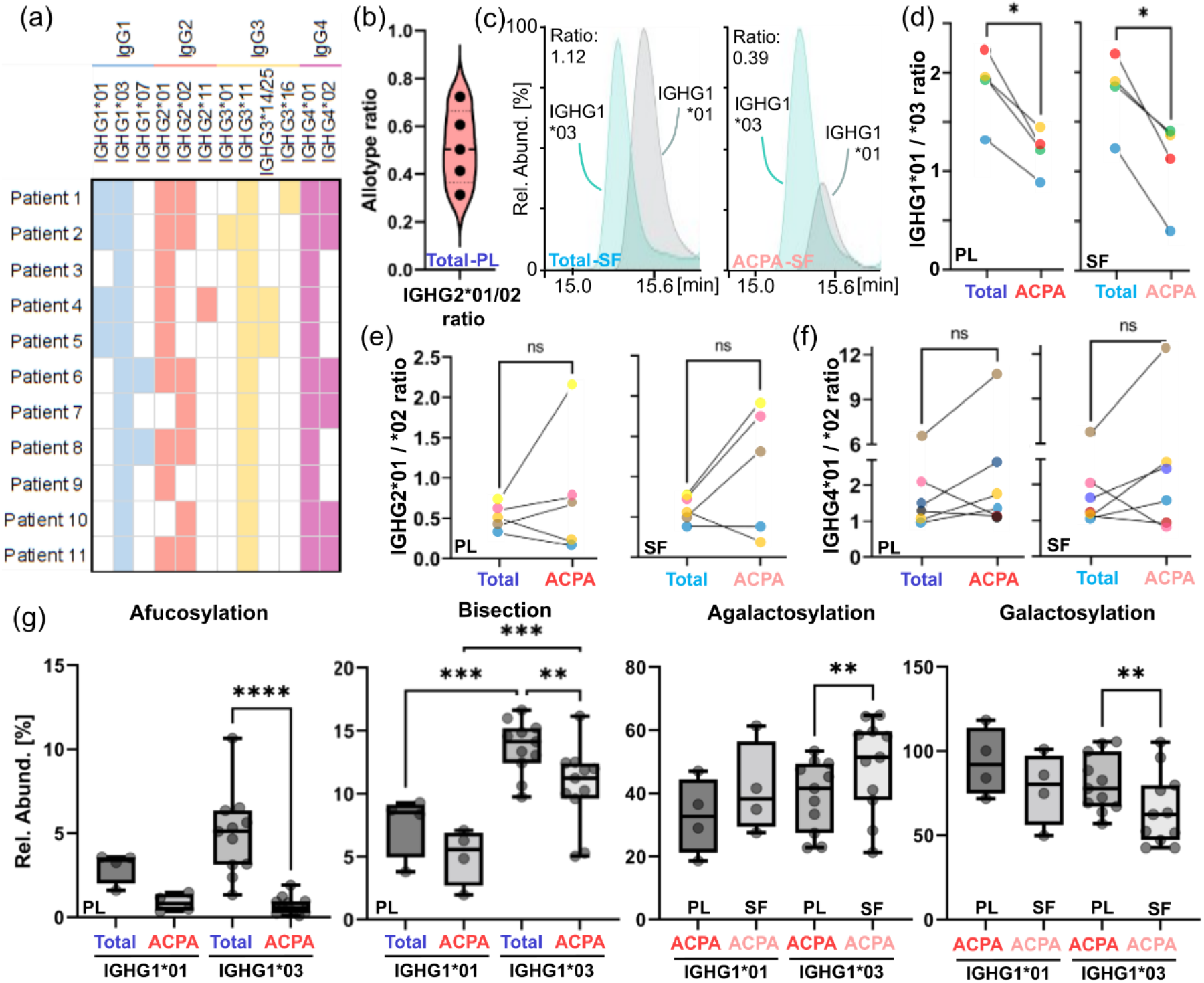
Allotype expression and allotype-specific glycosylation. (a) Allotype constitution of the patients studied. (b) Distribution of ratios of the IGHG2*01 and IGHG2*02 allotype in applicable patients for total plasma (PL) fractions. (c) EICs of the G0F glycoform of IGHG1*01 and IGHG1*03 allotypes for an exemplary patient (#4). The left panel exemplarily shows the allotype ratio of total IgG in SF, whereas the right panel demonstrates the shifted allotype ratio in ACPA from SF. Allotype ratios (considering all glycoforms) of (d) IGHG1*01/03, (e) IGHG2*01/02, and (f) IGHG4*01/02, respectively, in total and ACPA IgG in plasma and SF of applicable patients. Individual patients were color-coded throughout panels d-f. Significance levels were assessed by paired t-tests. (g) Allotype-specific derived traits in total and ACPA IgG from plasma and SF. Individual patients are indicated. Mixed-effects analysis considering paired values followed by Šídák’s multiple comparisons test was employed (* for p ≤ 0.05, ** for p ≤ 0.01, *** for p ≤ 0.001, and **** for p ≤ 0.0001).

Fc subunit profiling allowed allotype-specific glycosylation analysis of all subclasses. Allotype-dependent glycosylation changes were investigated in detail for IgG1 allotypes (IGHG1*01 and IGHG1*03) (**Figure 3g**). Although systematic differences in glycan traits afucosylation and bisection were observed between these allotypes, the alterations between total and ACPA IgG were conserved for both IgG1 allotypes. The same was true for the comparison of glycan traits galactosylation and agalactosylation between ACPA IgG from plasma and SF. Systematic glycosylation differences were specifically pronounced for bisection comparing IGHG1*01 and IGHG1*03 allotypes.

### IgG3 C_H_3 domain glycosylation and occupancy is altered in ACPA fractions

Depending on the allotype, IgG3 may bear a second *N*-glycosylation site in its C_H_3 domain. For the first time, we report on the presence and variability of these IgG3 C_H_3 domain glycans in total IgG and ACPA of RA patients (**Figure 4**). Since both glycosylation sites are present on the Fc subunit and the C_H_2 domain glycosylation site is virtually fully occupied, glycosylation of the C_H_3 domain was indicated by the presence of a doubly glycosylated IgG3 Fc subunit at higher masses. C_H_3 domain glycans were characterized in their interplay with C_H_2 domain glycans but also without the influence of C_H_2 domain glycans by the utilization of the EndoS2 glycosidase. We demonstrated that EndoS2 was able to specifically and efficiently cleave the chitobiose core of C_H_2 domain glycans, yet leaving C_H_3 domain glycans unaltered. In **Figure S7**, we show the selective C_H_2 glycan cleavage for a monoclonal IgG3 in more detail. Exemplary doubly glycosylated IgG3 spectra are shown for total IgG (plasma) and ACPA IgG (SF) for the most common IgG3 allotype IGHG3*11 (**Figure 4a**). Site-specific glycosylation data on both C_H_2 domain (obtained from Fc/2 comprising a non-glycosylated C_H_3 domain) and C_H_3 domain glycans (obtained from EndoS2-treated C_H_3 domain glycosylated Fc/2) was integrated to assess their contribution to the doubly glycosylated spectrum (**Figure S8**). A complete annotation of the doubly glycosylated IgG3 spectrum is provided for total IgG (plasma) in **Table S4**. Interestingly, a significant increase in high mannose-bearing glycans was found in the C_H_3 domain of ACPA compared to total IgG (**Figure 4b**). This difference was consistent for both plasma and SF. This finding could be confirmed after deglycosylation of the C_H_2 domain (**Figure S9**). Generally, the occupancy of the C_H_3 domain glycosylation site of IgG3 (IGHG3*11) was found to be relatively variable between patients, which was ranging from 7.2% (ACPA SF; patient 11) to 45.1% (ACPA plasma; patient 3). Moreover, this proportion of occupancy was altered between total and ACPA IgG, even though it did not reach significance. Next, glycosylation traits were assessed for doubly glycosylated IgG3 (**Figure 4c**): The significant differences reported were in line with the observations made for C_H_2 domain glycans in IgG3 as depicted in **Figure S5** and **S6**. In essence, the biofluid (plasma or SF) from which ACPA originated may be discriminated based on (a)galactosylation and sialylation levels, whereas total and ACPA IgG differed mainly in bisection. Fucosylation was not assessed as C_H_3 domain glycans showed virtually no core fucosylation (**Figure S9**). To exclude a potential bias in the Fc-based capturing of certain doubly glycosylated IgG3 glycoforms compared to antigen-specific capturing, we assessed this for monoclonal IgG3 (**Figure S10**). No evidence for a bias in capturing approaches was found.

**Figure 4.**
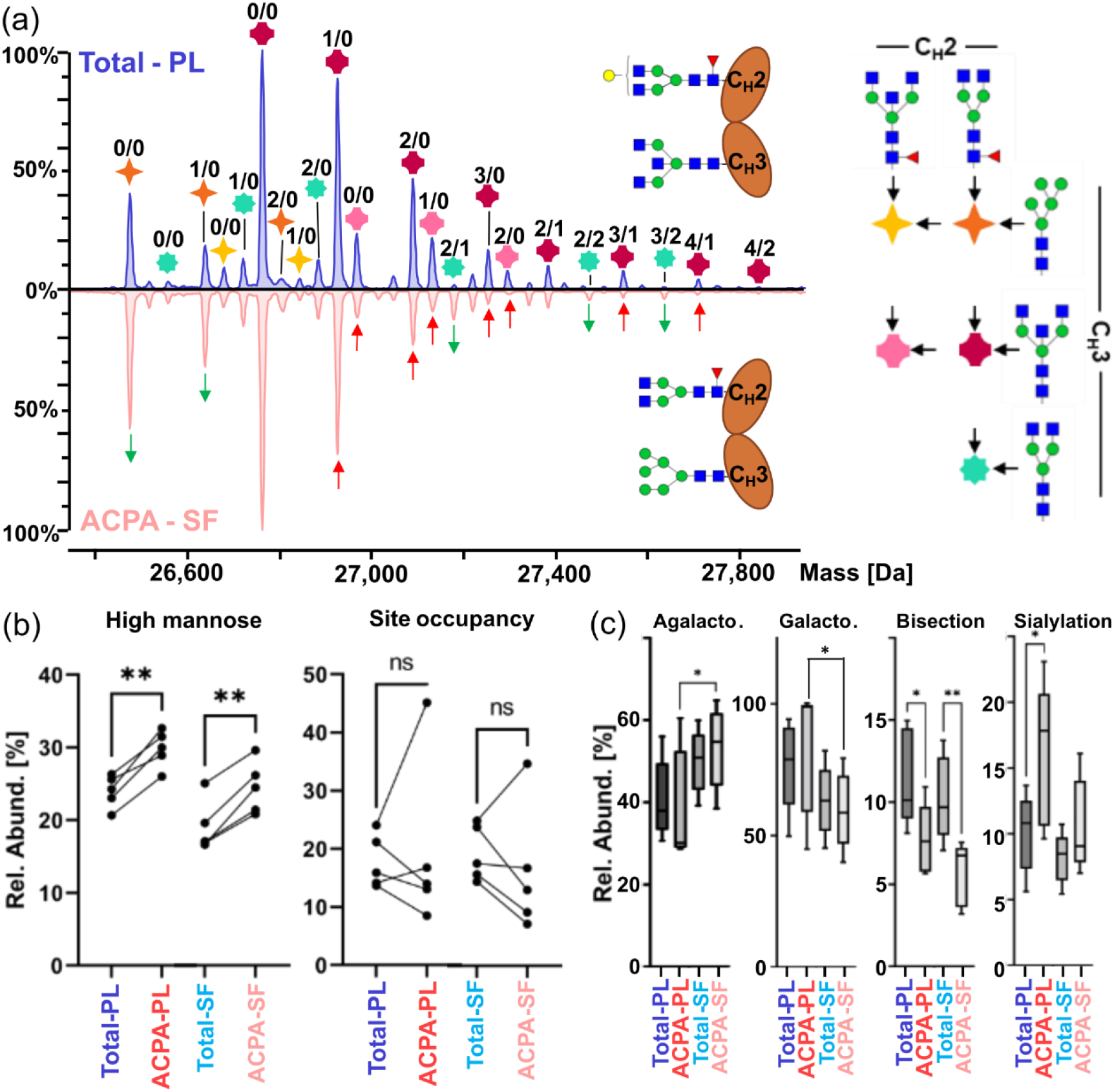
Assessment of C_H_3 domain glycosylation in IgG3. (a) Representative mass spectra of doubly glycosylated IgG3 (IGHG3*11 allotype) in total IgG from plasma (PL; blue) and in ACPA IgG from SF (light red). Observed masses are combinations of two *N*-glycans and annotated by symbols as shown in the legend on the right. These non-galactosylated and non-sialylated base structures are elongated by hexoses and sialic acids as indicated by the respective numbers above the symbols (the number of hexoses is listed as a first digit and the number of sialic acids as second digit). Differences between the two mass spectra were indicated by arrows (green arrows indicate a higher abundance and red arrows a lower abundance of a glycan peak in ACPA IgG (SF) compared to total IgG (plasma)). (b) The abundance of high mannose-bearing doubly glycosylated IgG3 as well as the glycosylation site occupancy of the C_H_3 domain of IgG3 were quantified in the different IgG fractions. Paired t-tests were applied. (c) Derived traits, i.e., agalactosylation, galactosylation, sialylation, and bisection, were assessed for the doubly glycosylated IgG3. Of note, these traits were influenced by both C_H_2 and C_H_3 domain glycans. One-way ANOVA considering paired values followed by Šídák’s multiple comparisons test was employed to assess significance (* for p ≤ 0.05, and ** for p ≤ 0.01).

### Fc subunit characterization reveals additional IgG proteoforms

Next to allotypes, subclasses, and glycosylation, information on additional post-translational modifications (PTMs) of the Fc could be retrieved. An overview of identified and monitored PTMs is depicted in **Figure 5a**. Incomplete lysine clipping was exclusively found in total and ACPA IgG purified from SF (**Figure 5b**). Significantly more C-terminal lysine was found in ACPA IgG1 and IgG4 compared to total IgG1 and IgG4. For a subgroup of patients, C-terminal GK truncation was observed in IgG4 (**Figure 5c**). This was found to be lower in SF compared to plasma for total IgG. Due to the generally lower abundance of ACPA compared to total IgG, this feature could only be assessed in a few patients for ACPA. In addition, disulfide bond formation was found to be incomplete in the Fc as indicated by identification of a N-terminally truncated variant (**Figure 5d**) as well as cysteinylation (**Figure 5e**). Fc/2 portions comprising reduced disulfides could be chromatographically separated and showed a mass shift of approx. +2 Da as additionally demonstrated for a monoclonal IgG1 antibody (**Figure S11**). Reduction or alkylation led to a loss of these distinct peaks, supporting the proposed identity of these peaks. Although deamidation may potentially lead to a similar mass shift, forced deamidation did not affect the abundance of these late eluting proteoforms (**Figure S11**). In total and ACPA IgG, late eluting proteoforms showing +2 Da mass shifts were also observed for IgG4 allotypes. All quantified molecular features of the Fc in the different IgG fractions were assessed by another more comprehensive PCA (**Figure S12**). Inclusion of additional parameters, e.g., allotype ratios, IgG3 C_H_3 domain glycosylation, and lysine variants, further enhanced the separation of analyzed IgG fractions. In fact, the separation of total IgG from plasma and SF became apparent, besides an even clearer distinction between ACPA from plasma and SF.

**Figure 5.**
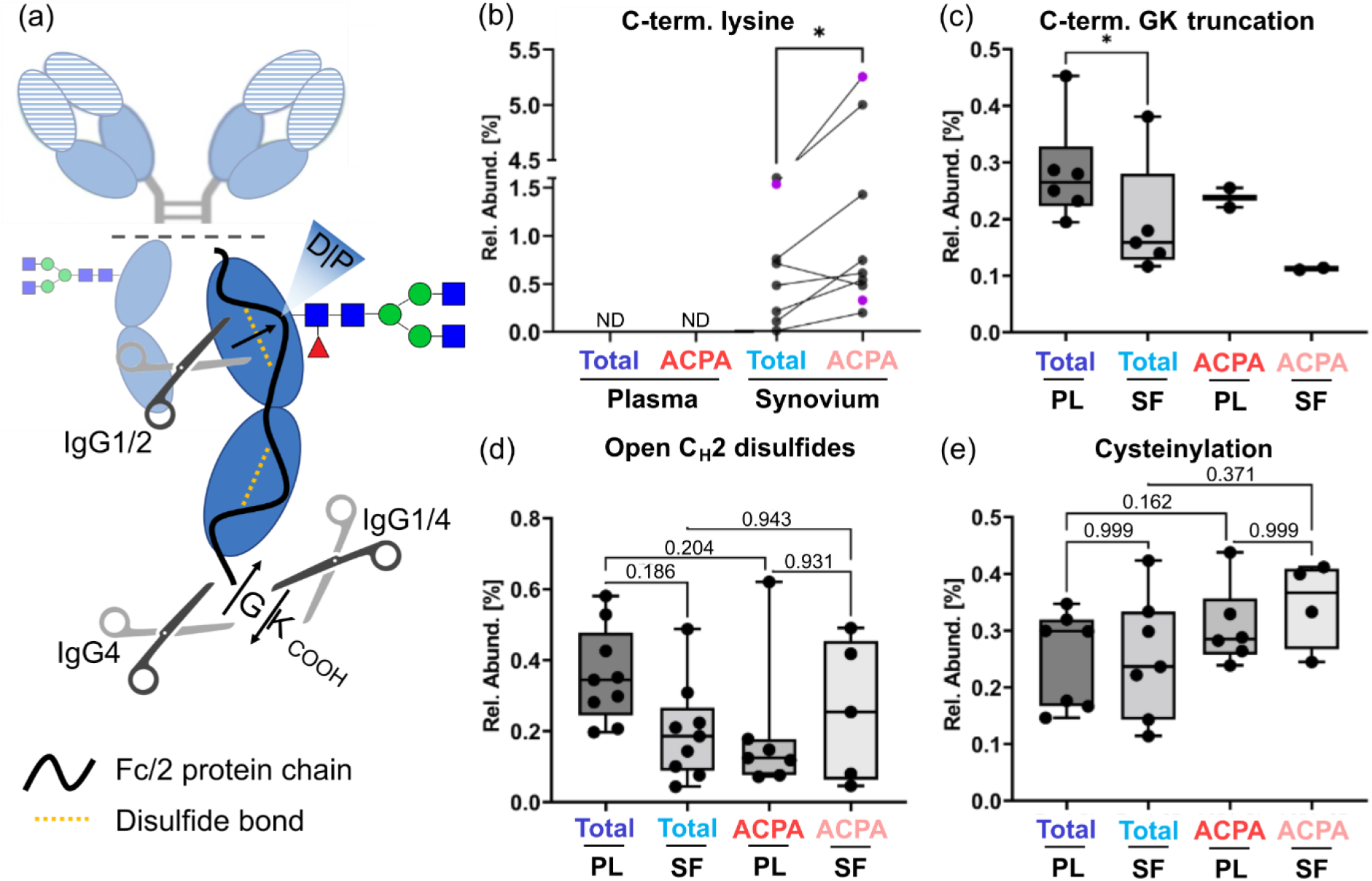
Additional PTMs identified in the Fc region. (a) Schematic of the Fc/2 portion, which visualizes the location and nature of observable PTMs, namely incomplete C-terminal lysine clipping, C-terminal GK truncation, open C_H_2 disulfides, and cysteinylation. (b) Relative abundance of C-terminal lysine in IgG1 (black) and IgG4 (violet) purified from SF. Notably, IgG from plasma had no detectable levels of C-terminal lysine. A paired t-test was employed. Box-whisker plots (minimum to maximum; including individual values) of (c) C-terminal GK truncation of IgG4, (d) open C_H_2 disulfides, and (e) cysteinylation. Of note, open C_H_2 domain disulfides could be assessed specifically due to the observable hydrolysis between amino acids DP, which are located between the two C_H_2 cysteines. Simultaneous DP hydrolysis and open C_H_2 disulfides lead to the loss of a relatively big N-terminal amino acid stretch. The resulting truncated Fc/2 was used for relative quantification. Statistical significance was assessed by either (c) paired t-testing or (d, e) mixed-effects analysis followed by Šídák’s multiple comparisons test (* for p ≤ 0.05). For panels (d) and (e) adjusted p-values are indicated.

### Fc proteoforms associate with markers of disease activity

Identified IgG features were screened for correlations with ACPA IgG levels and clinical parameters, i.e., erythrocyte sedimentation rate (ESR) and disease activity score 28 (DAS28). On the one hand, a PCA was compiled that relied on subclass abundances and subclass-specific glycan traits from all four investigated IgG fractions leading to a separation of patients according to the variation in these characteristics (**Figure 6a**). Overlaying clinical parameters revealed that both DAS28 as well as ESR were elevated in patients that were low in principal component 1 (PC1). Although PC1 is a linear combination of multiple variables, galactosylation next to non-glycosylated IgG levels were a strong contributor to PC1 values (**Figure S13**). No apparent association of ACPA IgG levels was found in this PCA analysis. On the other hand, pairwise correlation analysis was performed for IgG features separated into the four IgG fractions (**Figure 6b** and **c**). Regarding glycosylation, ESR and DAS28 were found to be significantly positively correlated with agalactosylation of total IgG1 (plasma and SF). IgG1 galactosylation and sialylation were negatively correlated with ESR and DAS28 in total IgG but did not reach significance in ACPA IgG. Similar trends of correlation were observed for other subclasses in total IgG and also ACPA IgG but were not significant in most cases. Interestingly, also IgG4 abundances in total IgG were found to correlate with DAS28 and ESR.

**Figure 6.**
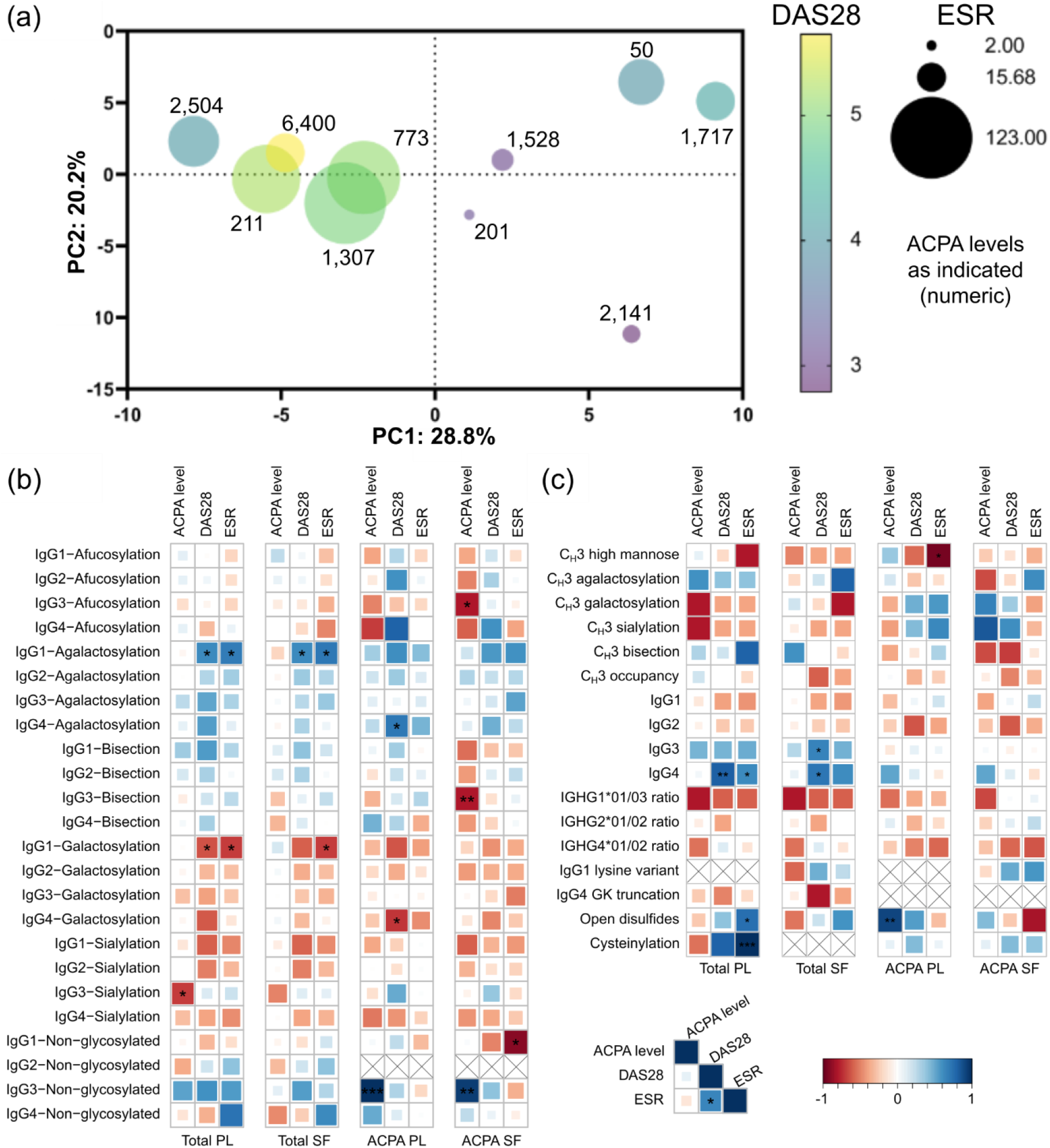
Associations of clinical parameters with IgG features. (a) PCA based on subclass abundances and subclass-specific glycosylation features summarizing all four investigated IgG fractions per patient. Circles represent individual patients for which clinical parameters where available. Size of circles indicates the ESR values, whereas the color represents the DAS28 score. ACPA IgG levels are indicated as numeric next to the circles. The corresponding loading plot is depicted in Figure S13. Of note, additional PTMs were excluded in this visualization due to the relatively higher proportion of missing values. (b) Correlation of glycosylation traits with available clinical parameters (ACPA IgG level, DAS28, and ESR) grouped by IgG fractions. (c) Correlation of additional IgG features and subclass abundances with aforementioned clinical parameters. In all cases Spearman correlation coefficients were calculated and indicated by size and colors of the squares (as shown in the legend). Significance levels are indicated (* for p ≤ 0.05, ** for p ≤ 0.01, and *** for p ≤ 0.001). Pairwise depletion was implemented for both correlation coefficient and p-value calculations. Features that were absent or based on less than four pairs in specific IgG fractions are marked with crosses in the respective squares.

## Discussion

Anti-citrullinated protein antibodies (ACPA) are highly specific biomarkers for RA providing diagnostic and prognostic information guiding clinical decision making. Determination of ACPA-associated variability, including subclass and allotype distribution in combination with PTMs such as glycosylation, may further expand insights into disease course and activity. However, current approaches are mostly centered on IgG1 Fc glycosylation potentially leading to a loss or neglect of crucial information. To close this gap, we employed a novel analytical approach based on the characterization of the intact Fc/2 subunit to comprehensively assess the wealth of ACPA IgG glyco- and proteoforms. ACPA and total IgG were profiled in paired samples of plasma and SF in a cohort of established RA patients providing insights into IgG fraction-specific molecular features and unraveling their correlation with disease.

Expanding on a recently established methodology, we comprehensively characterized Fc/2 subunits from total and ACPA IgG that were affinity-purified from plasma and synovial fluid of RA patients (**Figure 1a**) [39]. In contrast to (glyco)peptide analysis [52], the employed subunit-based methodology enabled us to specifically assess IgG allotypes and subclasses avoiding isomeric and identical (glyco)peptides as analytes [39]. Consequently, all Fc-associated PTMs including glycosylation were characterized in an allotype- and therefore subclass-specific manner expanding on current methodologies and providing novel insights into (auto)antibody characteristics [30]. Integrating this wealth of identified IgG features in a multivariate PCA (**Figure 2a, Figure S12**) revealed highly uniform alterations across all patients dependent on the investigated fractions, i.e., total IgG (plasma/SF) and ACPA IgG (plasma/SF). Together this resulted in a clear distinction in the PCA. However, already on the level of subclass distributions we observed substantially patient-dependent autoimmune responses that were additionally differing between plasma and SF. As reported earlier [30, 35, 36], relative abundances of IgG1 and IgG4 were elevated in ACPA while those of IgG2 and IgG3 were decreased compared to total IgG (**Table S3**). The difference of the IgG1/4 ratio between total and ACPA IgG differed for the individual patients and indicated that some ACPA responses were almost exclusively IgG1-driven, while others show a strong contribution of IgG4 (**Figure 1b**). Strikingly, for some patients we found high levels of IgG4 even at the level of total IgG. Total IgG4 but not ACPA IgG4 levels were positively correlating with disease activity-related parameters, namely DAS28 and ESR (**Figure 6**). Given the fact that IgG4 are generally relatively poor activators of complement and Fcγ receptors (FcγRs) [51], the absence of a correlation in ACPA seems plausible. The finding that total IgG4 levels correlated with mentioned inflammatory markers is supported by a significant but relatively weak correlation of these two factors in a recent meta-study and is worth further investigation [53]. Next to subclasses, we assessed IgG sequence variations (allotypes) and monitored their relative abundance per subclass. Next to unequal expression of several allotypes in total IgG (**Figure 3b**), the ratios of IgG1, 2, and 4 allotypes differed between total and ACPA IgG for heterozygous patients (**Figure 3c, d, e**, and **f**). These ratios could only be compared between patients showing more common genetic combination of alleles, which reduced the number of applicable patients. Still, we could confirm the imbalance of IgG2 allotypes that was previously identified [54, 55]. Intriguingly, a significant difference in IgG1 allotype ratios was found between total IgG and ACPA both in plasma and SF (**Figure 3c**, and **d**). Potentially, this may arise from differences in the variable gene (V gene) repertoire linked to certain allotypes as suggested by Sarvas et al. [54]. Previously this group showed that the IgG2 allotype imbalance was dependent on the antigen used for vaccination against *Haemophilus influenzae* [54]. Generally, IgG allotype constitution may have functional implications in infectious [56] as well as autoimmune diseases [57]. The IgG allotype constitution of RA patients has been compared to healthy controls revealing little association [58] apart from an enrichment of certain allotypes in HLA-DR4-positive RA [59]. In contrast to allotype constitution, information on allotype expression remains scarce due to a lack of suitable analytical approaches. The allotype-sensitive methodology employed in this study facilitates the discovery of associations between allotype expression and disease as well as monitoring of allotype constitution without the need of DNA sequencing approaches.

Investigated IgG fractions demonstrated strikingly differing glycosylation patterns within patients (**Figure 2, S5**, and **S6**). ACPA IgG1 showed significantly lower levels of afucosylation than total IgG1 in both plasma and SF. This effect was less pronounced in IgG2 and absent in IgG4 (**Figure S5**). Notably, IgG3 afucosylation levels were dependent on the body fluid as SF-related fractions showed lower levels of afucosylation. Afucosylation levels of total IgG1 remained comparable to healthy individuals [60]. While Scherer et al. [33] did not find differences in ACPA IgG1 afucosylation, our observation is in line with other reports on ACPA glycosylation [19, 30]. The strong effect seen in our cohort could potentially be linked also to the cohort characteristics, which had in general high degree of inflammatory and disease markers (**Table S1**). Since afucosylation drastically increases FcγRIII-binding and in turn antibody-dependent cytotoxicity (ADCC), our findings point towards a lower potential of IgG1 ACPA to induce ADCC [61]. This effect may be further increased by the lower amount of galactosylation found in ACPA from SF compared to plasma. Hypergalactosylation increases both FcγR- and C1q-binding resulting in increased activation of effector pathways [61]. Differences in sialylation followed the same trends as galactosylation but were especially pronounced for IgG3 and IgG4. Moreover, significantly decreased bisection in ACPA compared to total IgG may further decrease affinity to FcγRIIIa [62]. This difference could be observed in all subclasses but was most pronounced in IgG1 and IgG2. Partly, this may be driven by the higher expression of certain IgG1 allotypes, i.e., IGHG1*03, in ACPA fractions and a systematically higher degree of bisection in these allotypes. Unlike afucosylation, bisection was even lower in ACPA IgG from SF compared to plasma. In general, the observed differences between ACPA from plasma and SF is conceivably linked to the high degree of locally produced ACPA and potentially a different microenvironment in the inflamed joints [63, 64]. Given the high abundance of ACPA in SF [65], the differences in glycosylation and the according influence on the capacity of these antibodies to activate the immune responses may be especially relevant for ACPA found in SF – the site of the disease-associated inflammation. Contrarily to ACPA comprising Fc glycans that potentially lower their effector potential, these glycan traits did not correlate with disease parameters in case of fucosylation and negatively correlated in case of galactosylation. Galactosylation-levels have been reported to correlate with the ESR in RA [33] and seem to be generally associated with inflammatory settings [66]. Additionally, we report on the reduction of non-glycosylated IgG in ACPA compared to total IgG. Fc glycosylation is not required for B cell activation but for efficient binding to FcγRs and C1q to elicit effector functions [67, 68]. The pronounced reduction of these immunologically “silent” IgG suggests a higher degree of functional IgG in the ACPA fractions, however given the generally low abundance of non-glycosylated species in total IgG fractions (approx. 0.3%) this may not be relevant for the course of the disease. Still, from an immunological viewpoint, the strong alterations of non-glycosylated IgG are intriguing and call for the exploration whether this finding is specific for ACPA or a general feature of highly reactive (auto)antibodies.

This study demonstrates the variability of the C_H_3 domain glycosylation of IgG3 in IgG fractions of RA patients (**Figure 4**). In general, all investigated patients showed a drastic difference in the nature of the glycans present on the C_H_2 or the C_H_3 domain (**Figure S8**). These findings corroborate the only previous report on C_H_3 domain glycosylation [69]. In addition, we report on the applicability of EndoS2 to selectively deglycosylate the C_H_2 domain glycosylation site under native conditions to assess the C_H_3 domain glycans without interference – a finding of interest for further studies on the functional implications of the C_H_3 domain glycans (**Figure S7**). Notably, high mannose type glycans were significantly elevated in both ACPA fractions (**Figure 4**). An elevated proportion of high mannose type glycans may indicate an increased production and turn-over rate of ACPA as compared to total IgG as high mannose type containing glycans are usually cleared by mannose receptors [70]. The alteration in high mannose type glycans was accompanied by a change in the site occupancy of the C_H_3 domain, although it did not reach significance. Assuming that pairing of IgG3 heavy chains is not dependent on the occupancy of the C_H_3 domain glycosylation site, the occupancy levels reported in this study (13.7-24.1% in Fc/2 of total IgG from plasma) would indicate that 25.5-42.4% of IgG3 at least carry one C_H_3 glycan in the intact molecule. In general, the combined C_H_2 and C_H_3 domain glycosylation traits galactosylation, bisection, and sialylation followed the trends observed for C_H_2 domain glycosylation (**Figure 4** and **S6**). Unfortunately, little is known about the functional implications of C_H_3 domain glycans. Although the respective amino acid sequence motive of the C_H_3 domain glycosylation site (Asn392) is not directly involved in FcγR binding [71], conformational changes may be of functional relevance. As the FcRn is directly interacting with the C_H_3 domain, changes in glycosylation may affect half-life of IgG3. Given the high potential of IgG3 to induce complement- and FcγR-dependent effector functions, the impact of C_H_3 domain glycosylation and its association with disease warrants further investigation [41].

Harnessing the strength of the employed methodology to determine all Fc modifications, we assessed PTMs beyond glycosylation. Biofluid-dependent differences included the incomplete clipping of the C-terminal lysine residue. Plasma-derived IgG did not show detectable levels of C-terminal lysine, however IgG from SF comprised levels up to 1.6% in total IgG and up to 5.3% in ACPA IgG. This is an interesting finding that points toward a local production of ACPA in the synovium as C-terminal lysine is rapidly processed by carboxypeptidases in serum [72]. Thus, SF may have lower carboxypeptidase activity or lower IgG half-life. Incomplete lysine clipping does likely not affect FcγR and FcRn binding but influences complement activation due to reduction of the hexamerization potential of IgG [73, 74]. Notably, elevated carboxypeptidase B activity in RA and osteoarthritis is associated with a reduction of inflammatory-mediated damage in the joints due to cleavage of proinflammatory peptides of the complement system [75, 76]. In line with these observations, we detect a positive correlation trend of C-terminal lysine abundance with inflammatory and disease markers (**Figure 6**). C-terminal lysine may thus act as a surrogate to assess basic carboxypeptidase activity in the synovium providing insights into inflammatory processes. Further, we confirm a recent report that found C-terminal glycine clipping in IgG4 and show that this affects both investigated IgG4 allotypes [77]. Abundance of this PTM was slightly lower in synovial IgG but we could not assess associations with ACPA due to the low amount of samples in which this modification was identified. Mechanistic and functional insights for this PTM are lacking. IgG comprised low levels of unpaired disulfides and related PTMs, e.g., cysteinylation (**Figure 5d** and **e**). Open disulfides were deeply characterized for IgG1 and there is strong evidence that reduced disulfides are also present in IgG4 but the findings may extend to the other subclasses. Abundances of open disulfides as well as cysteinylation were neither significantly different between total and ACPA IgG nor between ACPA IgG from plasma and SF. Despite this finding not being directly implicated in RA, it may be relevant for quality assessment of pharmaceutical monoclonal antibodies showing that this modification may be present in therapeutics as well as in endogenous IgG.

The outlined work represents the first study to comprehensively characterize Fc-associated differences in RA (auto)antibodies in an truly allotype- and subclass-specific manner. Assessment of total and ACPA IgG from paired samples of plasma and SF in a cohort of RA revealed intriguing differences in the PTM profiles of these four investigated IgG fractions. Drastic alterations in allotype and subclass profiles where accompanied by prominent differences in glycosylation, e.g., high fucosylation in ACPA IgG from plasma and SF and low galactosylation in ACPA IgG from SF. IgG features, which are commonly neglected or not accessible for conventional approaches, such as full subclass-specificity, allotype ratios, IgG3 C_H_3 domain glycosylation, non-glycosylated IgG or C-terminal lysine variants were assessed revealing in part strong associations with autoantibody fractions. Importantly, IgG proteoforms rather than ACPA IgG levels correlated with inflammatory markers such as ESR and DAS28 and could separate patients into distinct groups depending on their disease status. We are aware that the relatively small cohort size may induce over-fitting of the described data or may have hampered the discovery of further changes between IgG fractions or certain associations to turn statistically significant. Thus, future extensions to larger RA cohorts including a stratification by subtypes of the disease will further strengthen and expand the findings presented in this study. Yet, we believe that this study showcases the potential and benefits of IgG subunit analysis for the comprehensive characterization of (auto)antibodies and emphasizes the implementation of such methodologies in future RA studies and beyond. Convinced by the benefits of the presented study design and employed methodologies, we call for similar studies in other autoimmune diseases shedding light onto the highly variable Fc proteome of IgG.

## Acknowledgements

This work was funded by the LUMC fellowship 2020 (to E.D-V.) and the ERC GlycanSwitch grant (101071386; to M.W.). Steven W. de Taeye and Gestur Vidarsson are acknowledged for providing monoclonal IgG allotypes. Figures were partially created in BioRender.com.

## Author contributions

Constantin Blöchl: Conceptualization, Software, Formal analysis, Investigation, Data curation, Writing – original draft, Writing – review & editing, Visualization. Eva Maria Stork: Formal analysis, Data curation, Resources, Writing – review & editing. Hans Ulrich Scherer: Conceptualization, Investigation, Resources, Supervision, Writing – review & editing. Rene E.M. Toes: Conceptualization, Investigation, Resources, Supervision, Writing – review & editing. Manfred Wuhrer: Conceptualization, Investigation, Resources, Supervision, Funding acquisition, Writing – review & editing. Elena Domínguez-Vega: Conceptualization, Resources, Supervision, Project administration, Funding acquisition, Writing – review & editing. All authors agreed to the final version of the manuscript.

