## Supplementary figures for "Fc proteoforms of ACPA IgG discriminate autoimmune responses in plasma and synovial fluid of rheumatoid arthritis patients and associate with disease activity"

### Table of contents

Figure S1. ACPA capturing and specificity.

Figure S2. Assessment of variation in ACPA capture and allotype distribution.

Figure S3. Assessment of variation in ACPA glycosylation in IgG1 and IgG4 allotypes.

Figure S4. Assessment of variation in total IgG glycosylation in IgG1 and IgG4 allotypes.

Figure S5. Overview of subclass-specific glycosylation traits grouped by IgG fractions.

Figure S6. Overview of subclass-specific glycosylation traits grouped by IgG fractions.

Figure S7. Assessing selective deglycosylation of the $C_H3$ domain of IgG3.

Figure S8. Annotation of doubly glycosylated IgG3.

Figure S9. $C_H3$ domain glycosylation of IgG3.

Figure S10. Assessing a potential $C_H3$ domain glycoform bias for IgG3 due to Fc-based capturing.

Figure S11. Reduced disulfide bonds in the Fc subunit.

Figure S12. Comprehensive PCA including all investigated IgG features.

Figure S13. Loading plot corresponding to the score plot shown in Figure 6.

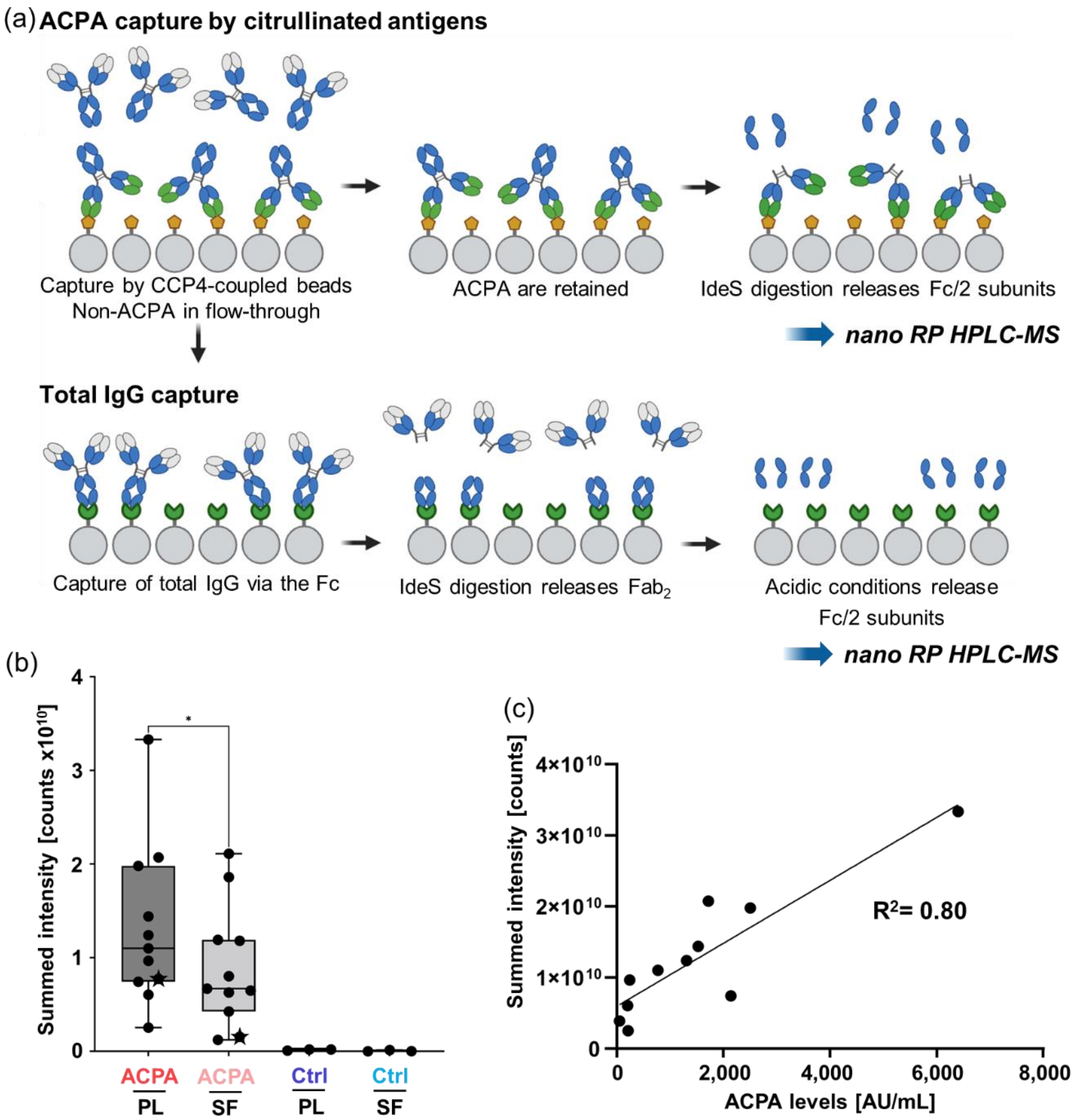


**Figure S1. ACPA capturing and specificity.** (a) Schematic outline of the capturing process. Plasma and SF samples were incubated with CCP4-coated beads that specifically capture ACPA. Non-ACPA were washed from the beads and were stored as flow-through. After several washing steps to reduce unspecific binding, IdeS was employed to cleave IgG below the hinge releasing Fc/2 subunits that were directly subjected to nano RP HPLC-MS analysis. Residual IgG ("Total IgG") were captured from the flow-through by Fc-specific beads and washed under neutral conditions. IdeS digestion released $Fab_2$ fragments that were subsequently washed from the beads retaining only Fc portions. Acidic conditions (0.1 M FA) were used to release Fc/2 subunits from the beads which were subjected also to nano RP HPLC-MS analysis without any prior purification steps. (b) Specificity of the capturing approach. The total ion current of an *m/z* region with high intensity for Fc/2 subunits (*m/z* 1180-1391) was used to approximate the total signal of Fc/2 subunits. This summed MS intensity was used to assess non-specific binding of the ACPA-capturing process. ACPA-positive samples (all investigated patients are depicted for plasma and SF) were compared to ACPA-negative samples (Ctrl). Of note, the total signals of patient 5 were indicated with stars in the plot since the applied volumes of sample were doubled due to low levels of ACPA detected in ELISA. Significance was assessed by a paired t-test (* for $p \leq 0.05$). (c) Summed MS intensities of ACPA IgG in plasma were linearly correlated with plasma ACPA IgG levels obtained by ELISA of the same patients. The two approaches showed a linear correlation with a $R^2$ coefficient of 0.80.

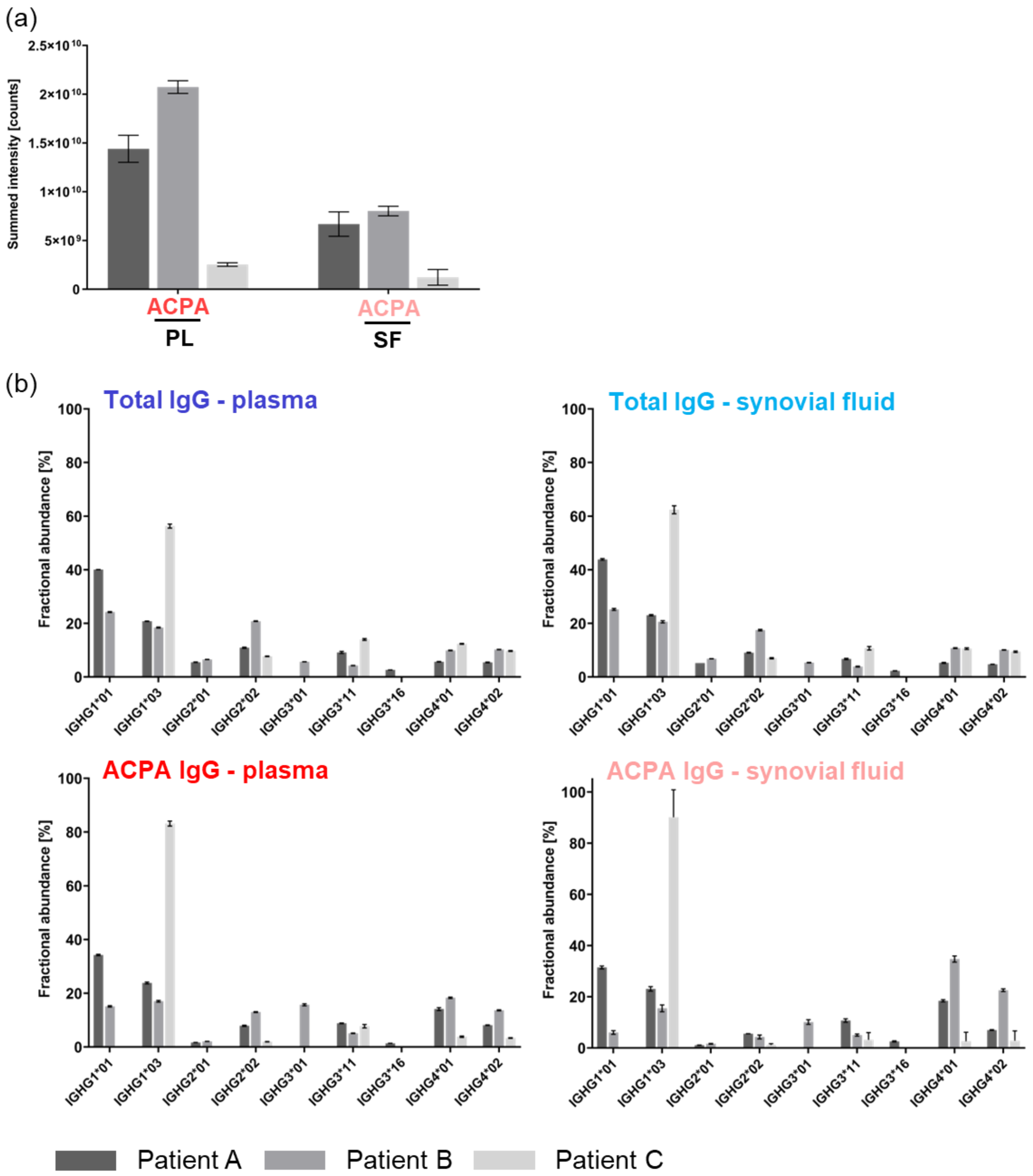


**Figure S2. Assessment of variation in ACPA capture and allotype distribution.** Patients A (# 1), B (#2), and C (#3) were investigated as three technical replicates including individual ACPA and total IgG capturing processes. Each of the triplicates was measured in duplicates by HPLC-MS. (a) Summed MS intensity (considering a *m/z* region of high intensity for Fc/2 subunits (*m/z* 1180-1391)) to assess ACPA capturing. Mean values and standard deviations were depicted for ACPA from plasma and SF. (b) Fractional allotype abundances in patients A-C determined for ACPA and total IgG purified from plasma and SF. Mean values and standard deviations were depicted.

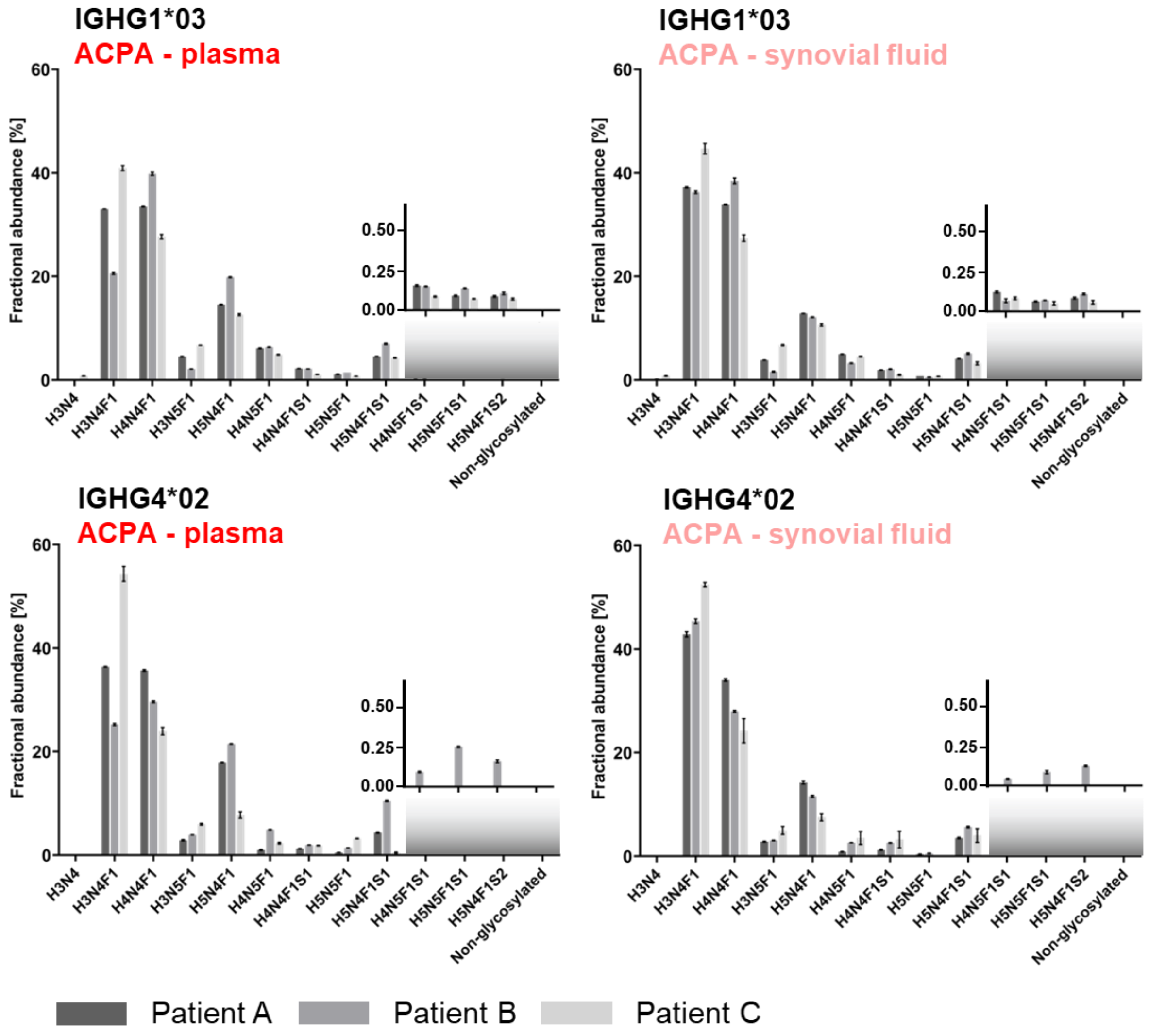


**Figure S3. Assessment of variation in ACPA glycosylation in IgG1 and IgG4 allotypes.** Glycosylation in highly prevalent IGHG1*03 and IGHG4*02 allotypes was characterized. Patients A (#1), B (#2), and C (#3) were investigated as three technical replicates including individual ACPA capturing processes. Each of the triplicates was measured in duplicates by HPLC-MS. Insets show a zoom (0.0-0.5%) into H4N5F1S1-, H5N5F1S1-, H5N4F1S2-carrying, and non-glycosylated Fc/2 subunits.

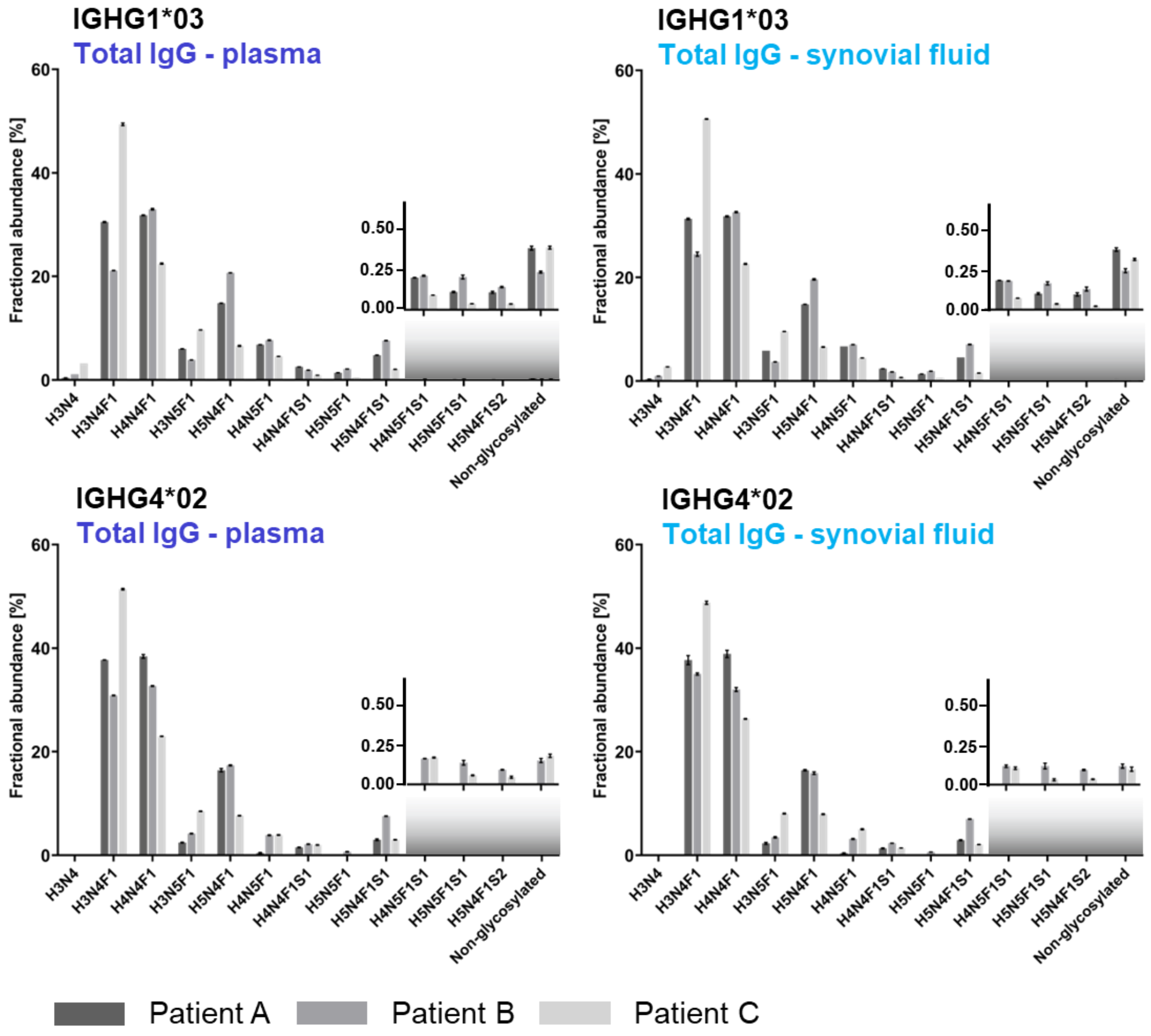


**Figure S4. Assessment of variation in total IgG glycosylation in IgG1 and IgG4 allotypes.** Glycosylation in highly prevalent IGHG1*03 and IGHG4*02 allotypes was characterized. Patients A (#1), B (#2), and C (#3) were investigated as three technical replicates including individual total IgG capturing processes. Each of the triplicates was measured in duplicates by HPLC-MS. Insets show a zoom (0.0-0.5%) into H4N5F1S1-, H5N5F1S1-, H5N4F1S2-carrying, and non-glycosylated Fc/2 subunits.

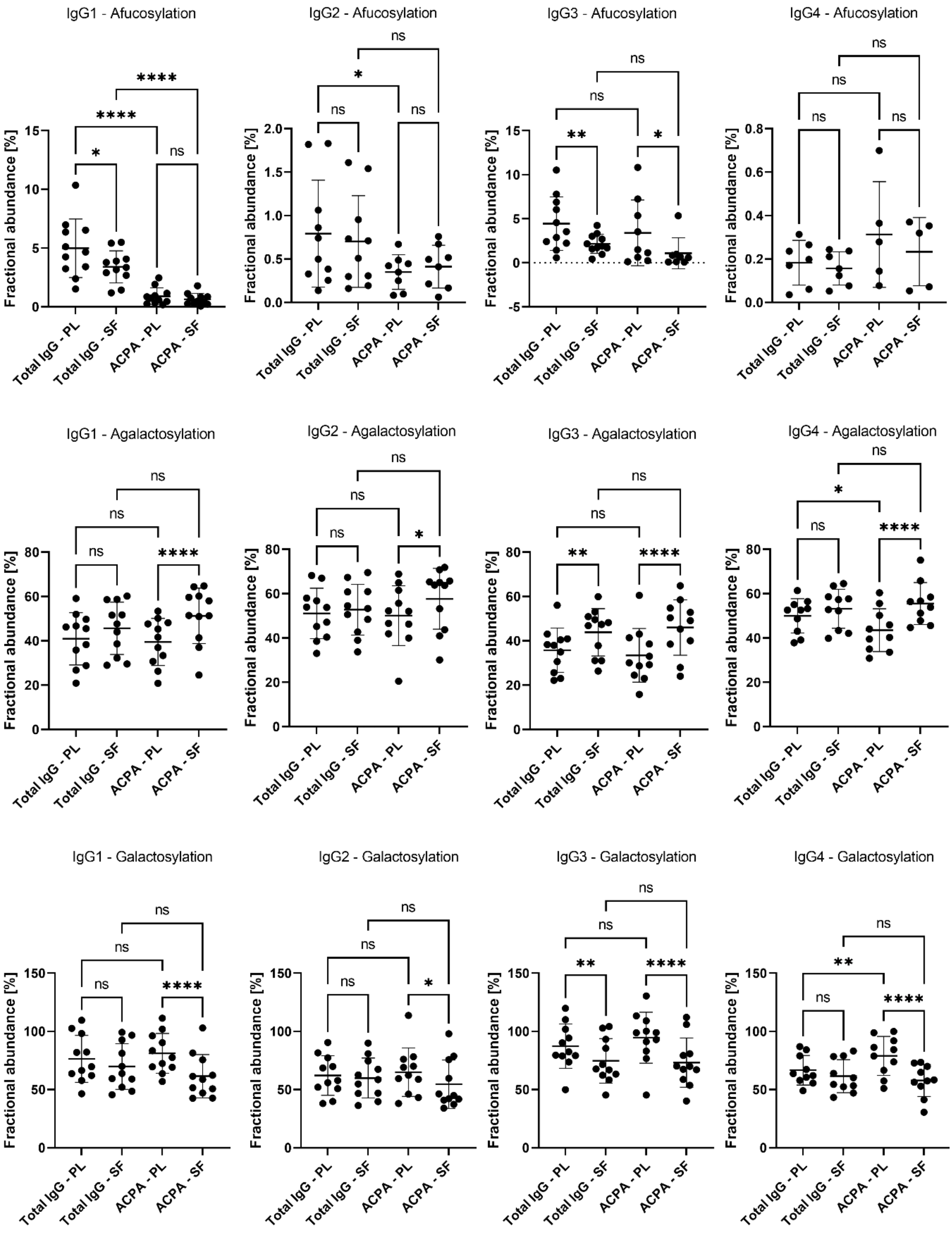


**Figure S5. Overview of subclass-specific glycosylation traits grouped by IgG fractions.** Afucosylation, agalactosylation, and galactosylation were compared between total and ACPA IgG from plasma (PL) and SF. Statistical significance was assessed considering paired observations by either one-way ANOVA or mixed-effects analysis if values were missing followed by Šídák's multiple comparisons test (predefined comparisons: total IgG plasma *vs.* total IgG SF; total IgG plasma *vs.* ACPA IgG plasma; total IgG SF *vs.* ACPA IgG SF; ACPA IgG plasma *vs.* ACPA IgG SF). Significance levels are indicated (* for $p \leq 0.05$, ** for $p \leq 0.01$, *** for $p \leq 0.001$, and **** for $p \leq 0.0001$).

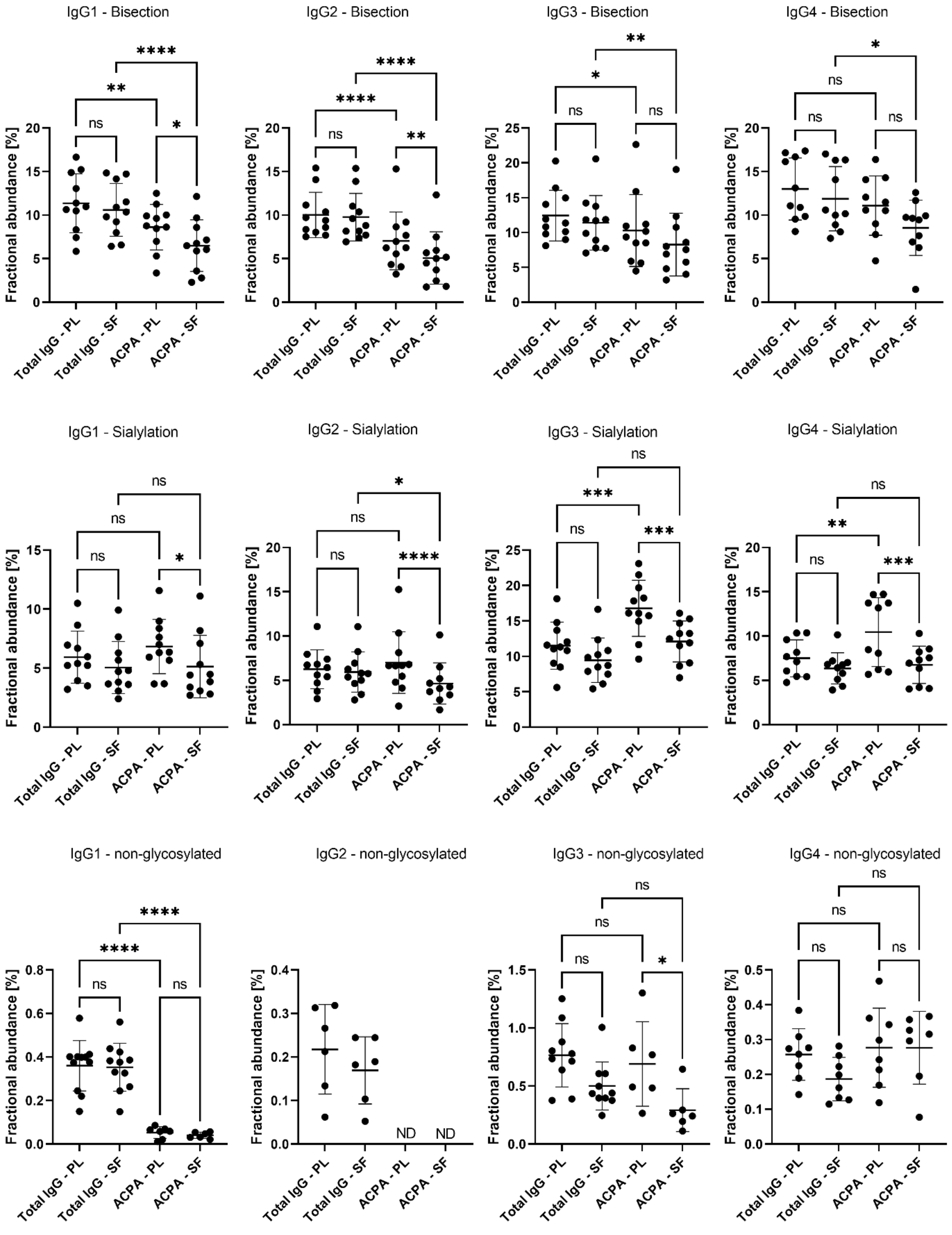


**Figure S6. Overview of subclass-specific glycosylation traits grouped by IgG fractions.** Bisection, sialylation, and non-glycosylated IgG were compared between total and ACPA IgG for plasma and SF. Statistical significance was assessed considering paired observations by either one-way ANOVA or mixed-effects analysis if values were missing followed by Šídák's multiple comparisons test (predefined comparisons: total IgG plasma *vs.* total IgG SF; total IgG plasma *vs.* ACPA IgG plasma; total IgG SF *vs.* ACPA IgG SF; ACPA IgG plasma *vs.* ACPA IgG SF). Significance levels are indicated (* for $p \leq 0.05$, ** for $p \leq 0.01$, *** for $p \leq 0.001$, and **** for $p \leq 0.0001$). "ND" indicates that the feature was detected in none of the patients in the specific fraction.

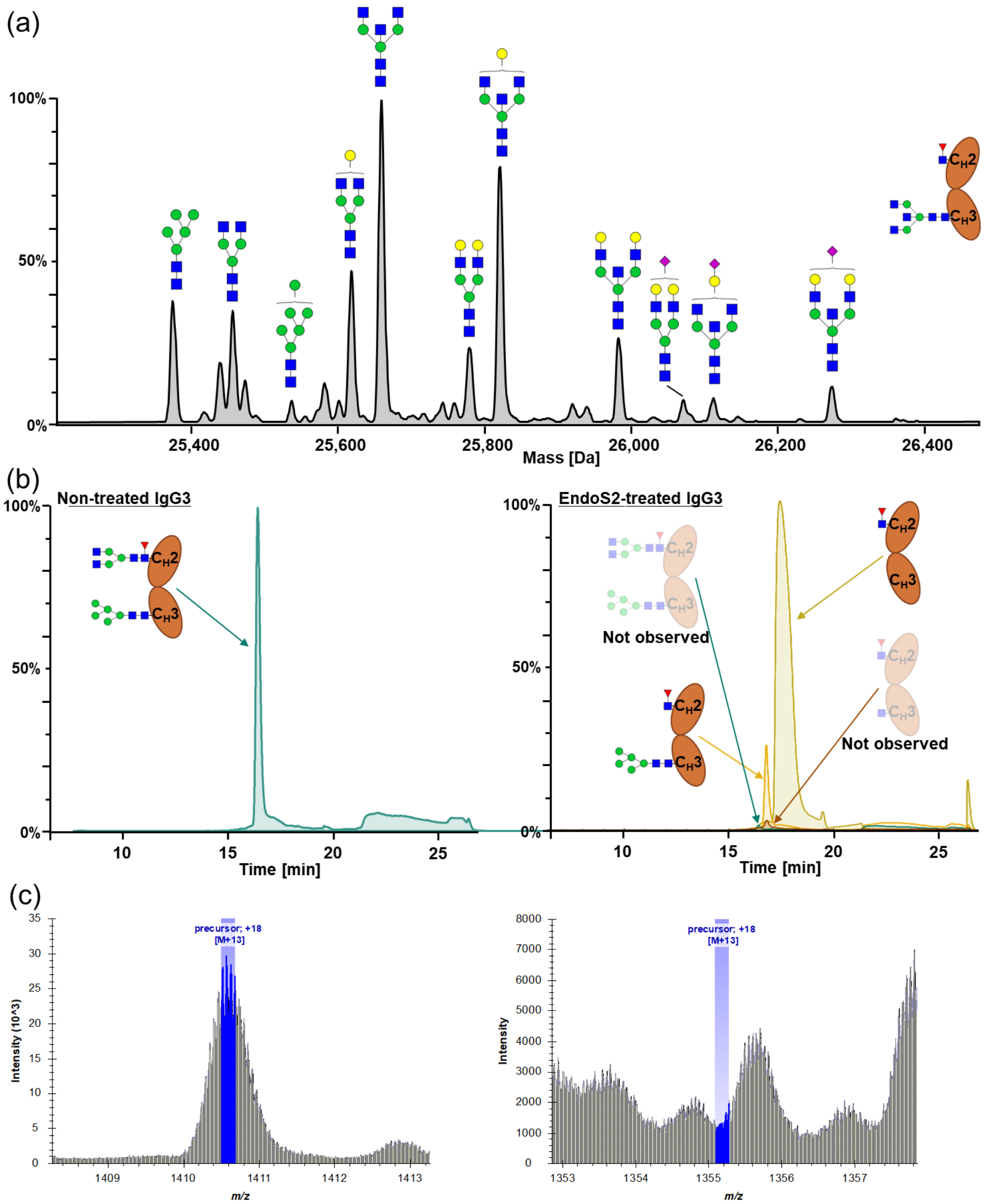


**Figure S7. Assessing selective deglycosylation of the $C_H3$ domain of IgG3.** $C_H3$ domain glycosylation in IgG3 could be site-specifically assessed after deglycosylation of the $C_H2$ domain glycans by the EndoS2 enzyme. (a) Exemplary mass spectrum obtained from total IgG of plasma from an RA patient. $C_H3$ domain glycans were annotated. (b) Specificity of EndoS2 cleavage was assessed in monoclonal IgG3 (IGHG3*11) before (left) and after EndoS2 treatment (right). To this end, four EICs were generated: Doubly glycosylated IgG3 (turquoise, H5N2/H3N4F1), $C_H2$ domain chitobiose core-cleaved IgG3 with occupied $C_H3$ domain glycosylation site (yellow, H5N2/N1F1), $C_H2$ domain chitobiose core-cleaved IgG3 without occupied $C_H3$ domain glycosylation site (dark yellow, N1F1), and doubly cleaved IgG3 (brown, N1/N1F1). As expected, out of the four investigated glycoforms only the doubly glycosylated IgG3 was detected for non-treated monoclonal IgG3. In contrast, $C_H2$ domain glycans were efficiently cleaved by EndoS2 as indicated by the two identified species after EndoS2 treatment. Neither non-cleaved nor doubly cleaved IgG3 was detected. (c) Corresponding mass spectra of the $C_H2$ domain chitobiose core-cleaved IgG3 with occupied $C_H3$ domain glycosylation site (left panel; mass in agreement) and the doubly chitobiose core-cleaved IgG3 (right panel; no evidence for this molecular species).

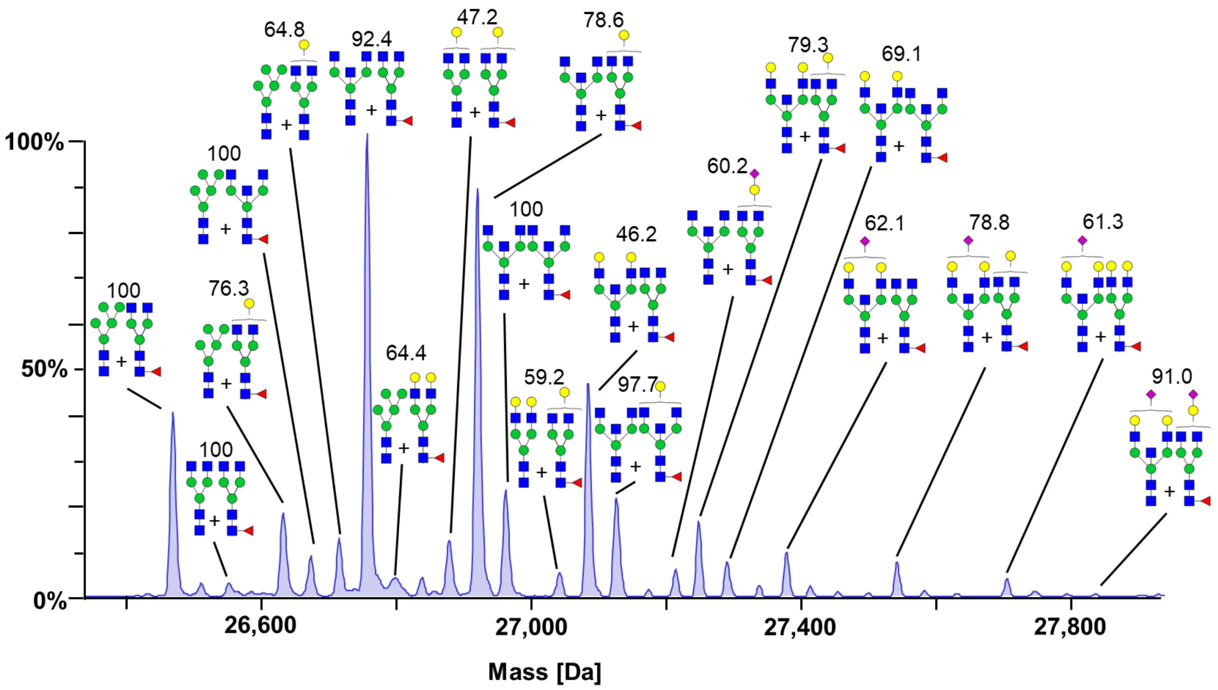


**Figure S8. Annotation of doubly glycosylated IgG3.** Quantitative site-specific information on $C_H2$ domain and $C_H3$ domain glycosylation was bioinformatically integrated employing the MoFi tool [1]. The most abundant combination of glycans for every peak in the mass spectrum was used for annotation. The left glycan is situated on the $C_H3$ domain and the right glycan on the $C_H2$ domain. The relative contribution (%) of the annotated glycoform to all possible glycoforms of each particular peak in the mass spectrum is shown on top of the glycoform. The most abundant mass peaks in the spectrum were annotated. A full list of annotations is provided in **Table S4**.

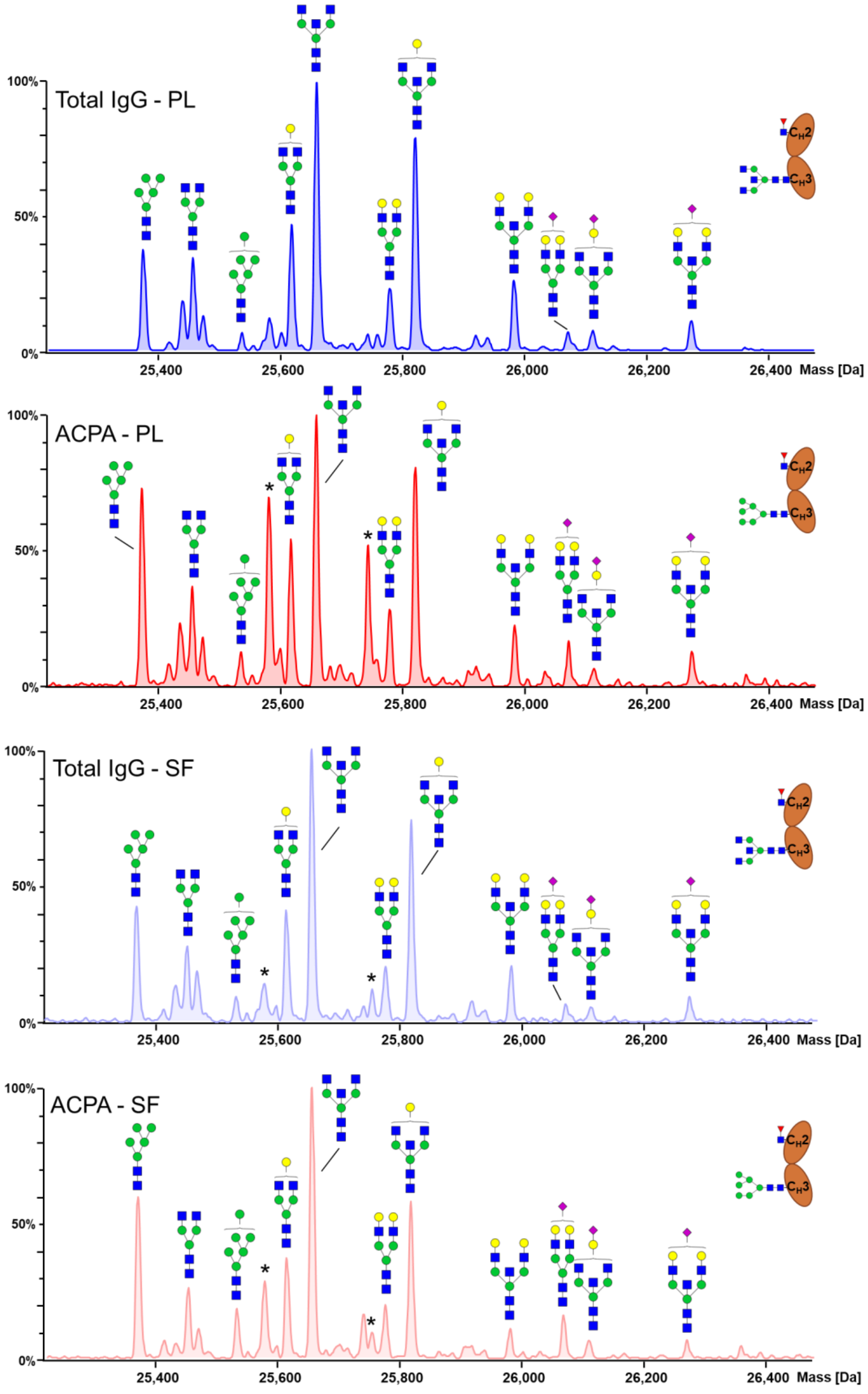


**Figure S9. $C_H3$ domain glycosylation of IgG3.** $C_H3$ domain glycosylation in IgG3 (IGHG3*11) was assessed after selective cleavage within the chitobiose core of $C_H2$ domain glycans with the endoglycosidase EndoS2. The four IgG fractions (total IgG and ACPA IgG from plasma and SF) of patient 3 were exemplarily shown. A strong shift in the relative abundance of high mannose type glycans was observed comparing total IgG to ACPA IgG (both plasma and SF). Asterisks mark co-deconvoluted masses related to IgG4.

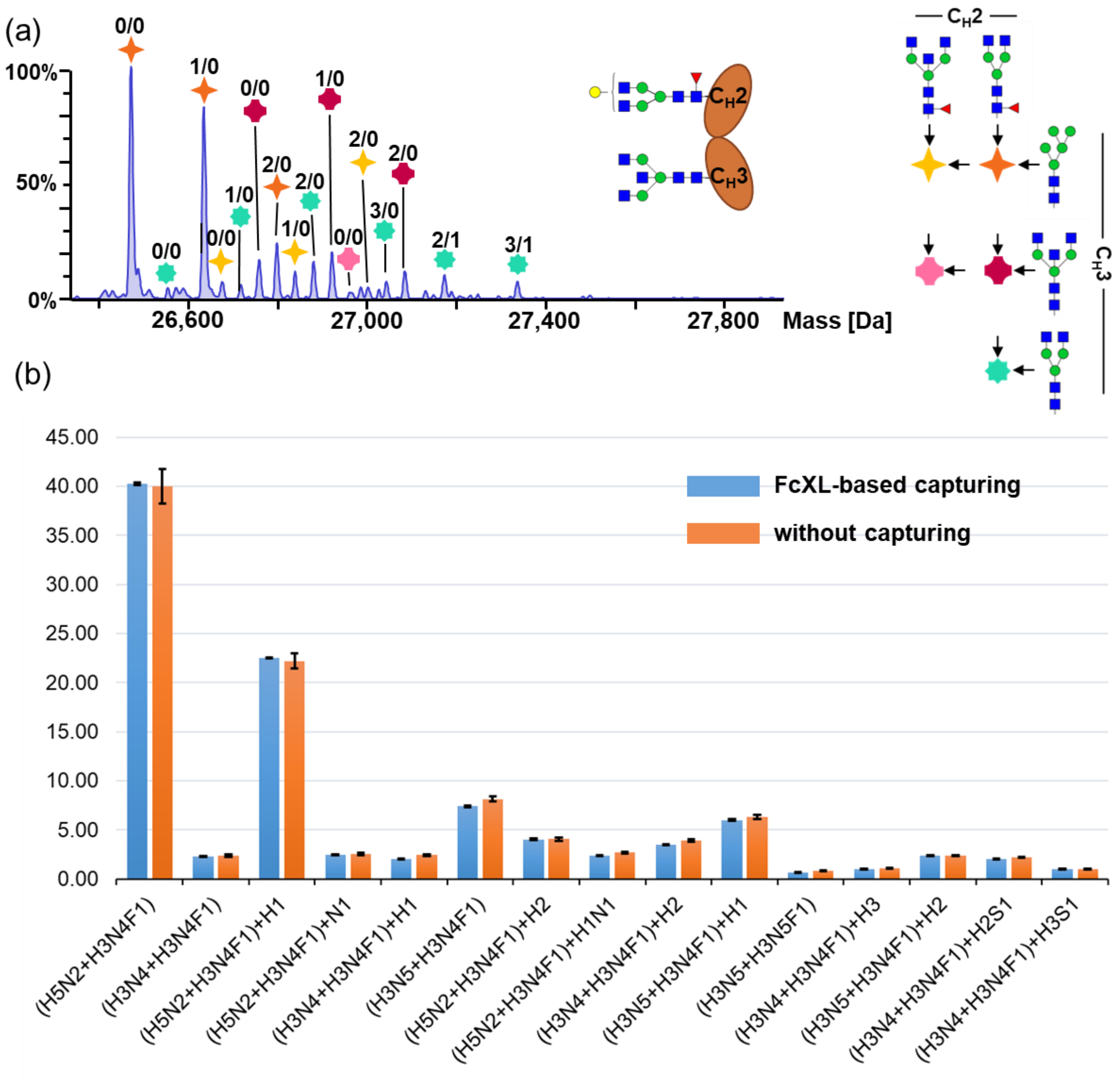


**Figure S10. Assessing a potential C$_H$3 domain glycoform bias for IgG3 due to Fc-based capturing.** (a) C$_H$3 domain glycosylation was characterized in a monoclonal IgG3 (IGHG3*11). As described in **Figure 4**, observed masses are combinations of two *N*-glycans and annotated by symbols as shown in the legend on the right. These non-galactosylated and non-sialylated base structures are elongated by hexoses and sialic acids as indicated by the respective numbers above the symbols (the number of hexoses is listed as a first digit and the number of sialic acids as second digit). (b) Identified doubly glycosylated glycoforms were relatively quantified with and without prior Fc-based capturing. Mean values and standard deviations are shown (n=3).

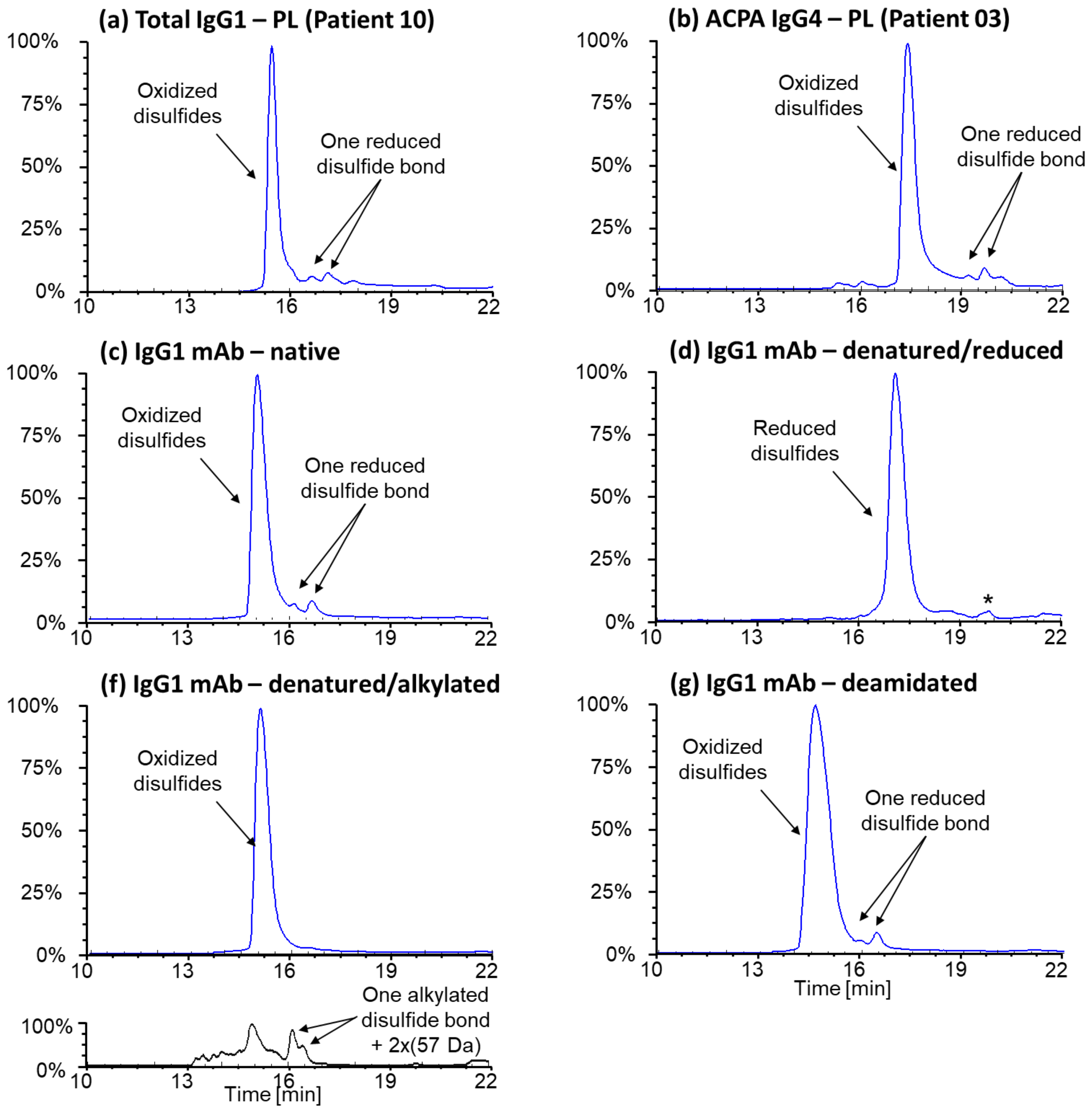


**Figure S11. Reduced disulfide bonds in the Fc subunit.** Late eluting peaks of low abundance were found in Fc/2 subunits obtained from patients, e.g. in IgG1 (a), and IgG4 (b) as well as from monoclonal IgG1 (c). A mass shift of approx. 1-2 Da indicated open intramolecular disulfide bonds. To confirm this, monoclonal IgG1 was (d) denatured and reduced leading to the disappearance of late eluting forms. The asterisk marks a co-extracted signal that does not show any relation to the Fc/2. Accordingly, (f) denaturation followed by alkylation with iodoacetamide resulted in the disappearance of these additional peaks paired with the appearance of two minor peaks showing two alkylated cysteines (+57 Da for carbamidomethylation). (g) For further corroboration, monoclonal IgG1 was extensively deamidated for 7 days under basic conditions to exclude that these peaks are caused by deamidated variants. Deamidation did not lead to an increase of these late eluting proteoforms. EICs of G0F glycoforms (19+) are shown in all cases. EICs were adapted for IgG4 and alkylated IgG1 accordingly.

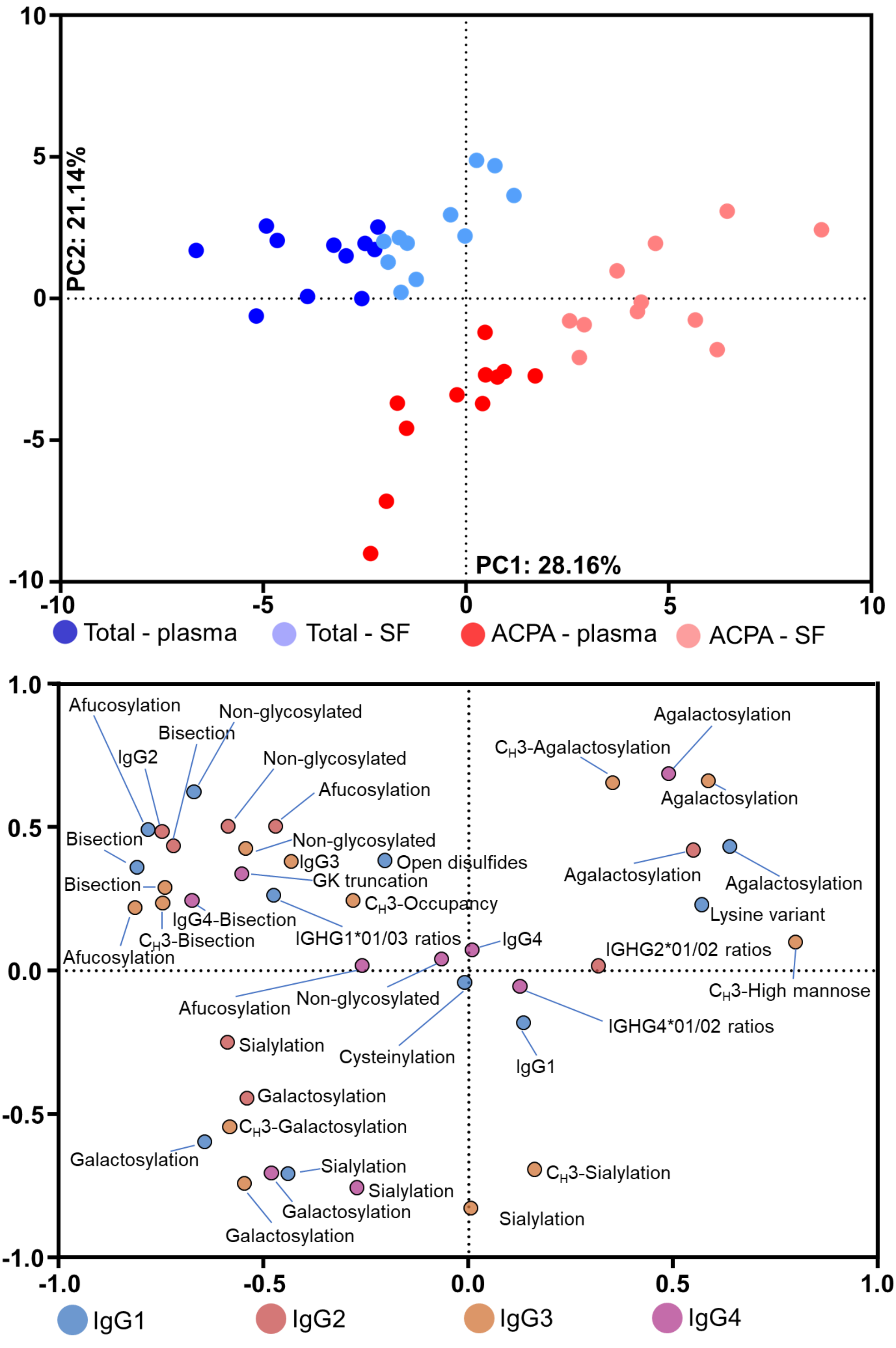


**Figure S12. Comprehensive PCA including all investigated IgG features.** Multivariate PCA was based on subclass abundances, subclass-specific glycosylation in the form of derived traits, IgG3 $C_H3$ domain glycosylation, allotype ratios, and additional IgG modifications. In the top panel, the score plot depicts data points representing individual patients. Each patient is represented by a set of four circles (blue: total IgG from plasma; light blue: total IgG from SF; red: ACPA IgG from plasma; light red: ACPA IgG from SF). In the bottom panel, the corresponding loading plot is shown. Variables were colored by their association with one of the four subclasses.

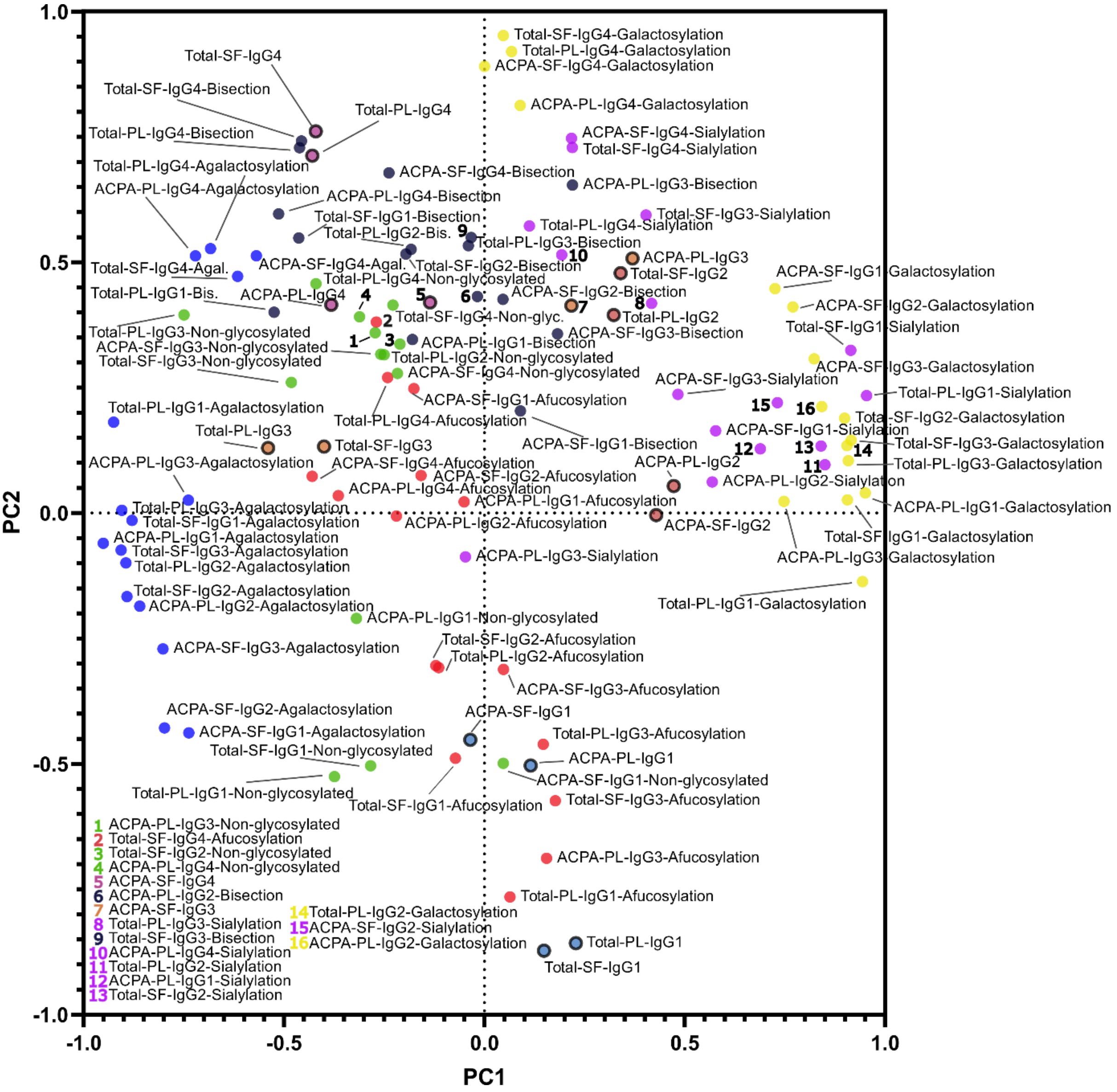


**Figure S13. Loading plot corresponding to the score plot shown in Figure 6.** Multivariate PCA was based on subclass abundances and subclass-specific glycosylation traits for all four investigated fractions, namely total IgG (plasma and SF) and ACPA IgG (plasma and SF). Included features and their respective contributions to PC1 and PC2 are depicted. Features were labeled accordingly if the space allowed to do so. If not, numbers were used to indicate IgG features. The legend for these numbers is included in the lower left side of the loading plot. Derived traits are color-coded: red (afucosylation), yellow (galactosylation), blue (agalactosylation), purple (sialylation), bisection (dark blue), and non-glycosylated (green). Subclass abundances are indicated with a thick black border line and color-coded: light-blue (IgG1), red (IgG2), orange (IgG3), and violet (IgG4).
